# Capturing the Conformational Heterogeneity of HSPB1 Chaperone Oligomers at Atomic Resolution

**DOI:** 10.1101/2024.12.16.628790

**Authors:** Raymond F. Berkeley, Alexander P. Plonski, Tien Phan, Kristof Grohe, Lukas Becker, Sebastian Wegner, Mark A. Herzik, Jeetain Mittal, Galia T. Debelouchina

## Abstract

Small heat shock proteins (sHSPs), including HSPB1, are essential regulators of cellular proteostasis that interact with unfolded and partially folded proteins to prevent aberrant misfolding and aggregation. These proteins fulfill a similar role in biological condensates, where they interact with intrinsically disordered proteins to modulate their liquid-liquid and liquid-to-solid phase transitions. Characterizing sHSP structure, dynamics, and client interactions is challenging due to their partially disordered nature, their tendency to form polydisperse oligomers, and their diverse range of clients. In this work, we leverage various biophysical methods, including fast ^1^H-based magic angle spinning (MAS) NMR spectroscopy, molecular dynamics (MD) simulations and modeling, to shed new light on the structure and dynamics of HSPB1 oligomers. Using split-intein mediated segmental labeling, we provide unambiguous evidence that in the oligomer context the N-terminal domain (NTD) of HSPB1 is rigid and adopts an ensemble of heterogenous conformations, the α-crystallin domain (ACD) forms dimers and experiences multiple distinct local environments, while the C-terminal domain (CTD) remains highly dynamic. Our computational models suggest that the NTDs participate in extensive NTD-NTD and NTD-ACD interactions and are sequestered within the oligomer interior. We further demonstrate that HSPB1 higher order oligomers disassemble into smaller oligomeric species in the presence of a client protein and that an accessible NTD is essential for HSPB1 partitioning into condensates and interactions with client proteins. Our integrated approach provides a high-resolution view of the complex oligomeric landscape of HSPB1 and sheds light on the elusive network of interactions that underly HSPB1 function in biological condensates.

**Significance statement:** HSPB1 is a ubiquitous cellular chaperone that helps prevent the aberrant aggregation of intrinsically disordered proteins involved in biological condensates and neurodegenerative diseases. Despite its central role in this process, many aspects of HSPB1’s structure and interactions with clients are not well understood due to its tendency to form polydisperse oligomeric structures and to function in heterogeneous condensate environments. Here, we present an integrated approach that includes segmental labeling, fast MAS NMR spectroscopy, and computational tools to characterize the structure and dynamics of HSPB1 in its oligomeric form and within client condensates at high resolution. Our approach, which is applicable to other complex biological systems, highlights the important role of HSPB1’s N-terminal domain in oligomeric assembly and interactions with clients.

## Introduction

Small heat shock proteins (sHSPs) are core regulators of proteostasis that bind a structurally diverse range of partially or completely unfolded protein clients (1–3). sHSPs also act in concert with ATP-dependent molecular chaperones to prevent misfolding or to prime misfolded or aggregated proteins for refolding (4). HSPB1 (also known as Hsp27) is an archetypal mammalian sHSP that interacts with a vast client network including many proteins linked to cancer and neurodegenerative diseases (5, 6). Mutations in HSPB1 have also been implicated in neurological disorders such as Charcot-Marie-Tooth disease and distal hereditary motor neuropathy (6). Similar to all sHSPs, HSPB1 exhibits a tripartite domain architecture comprising a central α-crystallin domain (ACD), flanked by an N-terminal domain (NTD), and a C-terminal domain (CTD), respectively (**Fig. S1a,b**) (7). Like other sHSPs, in the absence of a client protein, HSPB1 forms heterogenous and polydisperse oligomers, which range from dimers to large cage-like multimeric structures (8).

Biophysical and biochemical experiments have revealed detailed information regarding the structure, dynamics and interactions of HSPB1 dimers, which can be stabilized by mutations in the NTD and CTD (7). For example, it is well known that the ACD of HSPB1 consists of a β-sandwich fold (**Fig. S1b**) that is characteristic of all sHSPs (9–11). Two ACD domains come together to form the HSPB1 dimer, which results in the formation of a large six-stranded β-sheet. The dimer interface also forms a large groove (the dimer interface groove or the β3 groove) that can interact with the NTD and client proteins (12, 13). The dimer also has two identical edge grooves (also known as the β4/β8 grooves), where the side-chain of Ser155 forms a characteristic “bump” surrounded by two “holes” (7, 14). This bump-hole structure can interact specifically with the ^179^ITIPV^183^ motif in the CTD or segments of the NTD (12, 15). On the other hand, the NTD, which contains many proline and hydrophobic residues, is thought to sample an ensemble of “quasi-ordered” states that include interactions with the dimer and the edge grooves on the ACD, other NTDs and client proteins (12, 13, 15). Finally, the CTD is quite dynamic and it has been suggested that it can act as a solubility tag for the chaperone (7, 16).

While the dimeric HSPB1 form has been amenable to high-resolution structural analysis by X-ray crystallography and solution NMR spectroscopy, a comprehensive description of the HSPB1 oligomers has remained elusive due to their large size, structural heterogeneity and dynamics both in the client-free and client-engaged states (2, 3). Of note, solution NMR spectra of wild-type HSPB1 (i.e. in the absence of dimer-stabilizing mutations) contain signals only from the CTD, suggesting that this domain retains its fast dynamics in the oligomeric state (16). In the few structural models of sHSP oligomers that have been published to date, oligomers form cage-like structures composed of sHSP ACD dimers in various arrangements (17–20). In all of these cases, as well as HSPB1, the NTDs play an important role in the assembly and heterogeneity of the oligomers, a process that is also modulated by NTD phosphorylation (8, 12, 21). However, the exact nature of NTD interactions involved in HSPB1 oligomer assembly is currently unknown.

So far, the interactions of HSPB1 with three client proteins have been explored with solution NMR spectroscopy. This includes the Alzheimer’s disease related protein tau, which can form transient contacts with the NTD and specific contacts with the ACD edge grooves; however, only the interactions with the NTD appear to be productive (15, 22, 23). On the other hand, the low complexity domain of the stress response protein fused in sarcoma (FUS LC) can interact with the dimer groove on the ACD and the NTD depending on the context (13). HSPB1 has also been shown to modulate the behavior of FUS LC condensates, with the chaperone either disrupting liquid-liquid phase separation (LLPS) of FUS LC or preventing the aggregation of FUS LC depending on the oligomerization state of the chaperone (13). Finally, HSPB1 colocalizes with the TAR DNA-binding protein 43 (TDP-43) within stress granules (24) and appears to interact with the transient α-helical region in the low complexity domain of TDP-43 (25, 26). In all cases, however, the interactions could only be studied from the point of view of the client protein or in the context of mutation-stabilized HSPB1 dimers, while the point of view of the wild-type HSPB1 oligomers has remained inaccessible.

Here, we take advantage of recent advances in magic angle spinning (MAS) NMR spectroscopy, structural modeling, and molecular dynamics (MD) simulations to shed light on the complex and heterogeneous HSPB1 oligomers and characterize their structure and dynamics alone or in the presence of FUS LC condensates. To dissect the behavior of the tripartite domain architecture of HSPB1, we use split intein-mediated protein trans-splicing(27) to segmentally label the NTD and ACD-CTD domains of HSPB1 oligomers with ^13^C and ^15^N isotopes to generate an NMR dataset that reports unambiguously on the structure and dynamics of each domain. We also benefit from recent developments in MAS NMR probe development which allowed us to record high-resolution ^1^H-based multidimensional fast MAS NMR experiments (28) of fully protonated HSPB1 oligomer samples. Together with the details revealed by structural modeling with AlphaFold2 (29, 30) and atomistic and coarse-grained molecular dynamics (MD) simulations (31, 32), these tools have allowed us to describe the overall structure and dynamics of each HSPB1 domain in the oligomeric state, to capture distinct chemical environments in the ACD domain, and to shed light on the changes in oligomer size and domain dynamics in the presence of a phase-separated client protein. Our integrative approach, which is applicable to many other complex and heterogeneous biological systems, highlights the intricacy of interactions and states that define the biological function of the HSPB1 chaperone.

## Results

### HSPB1 forms cage-like polydisperse oligomers with heterogeneous architectures

Our first goal was to characterize the oligomerization landscape of wild-type recombinant HSPB1 in the client-free state (**Fig. S1c,d**). To this end, we employed mass photometry and analytical centrifugation experiments (AUC) at different concentrations (**Fig. 1a, b**). The formation of oligomers was concentration dependent, with dimers being the predominant species at low concentrations (250 nM), and tetramers and higher order oligomers present at higher concentrations (500 - 1000 nM) (**Fig. 1a, Fig. S2a**). In higher concentration samples (10 μM), AUC primarily detected 12-mers (**Fig. 1b**). The peak profiles remained the same in high and low salt conditions, and in the absence of reducing agent (**Fig. 1b, Fig. S2b,c**). Analysis by negative-stain transmission electron microscopy indicated the presence of particles that are 15 - 20 nm in diameter (**Fig. S3**). Samples were tested for activity using an insulin aggregation assay that showed decrease in aggregation as a function of increasing HSPB1 concentration (**Fig. S4**).

**Figure 1.**
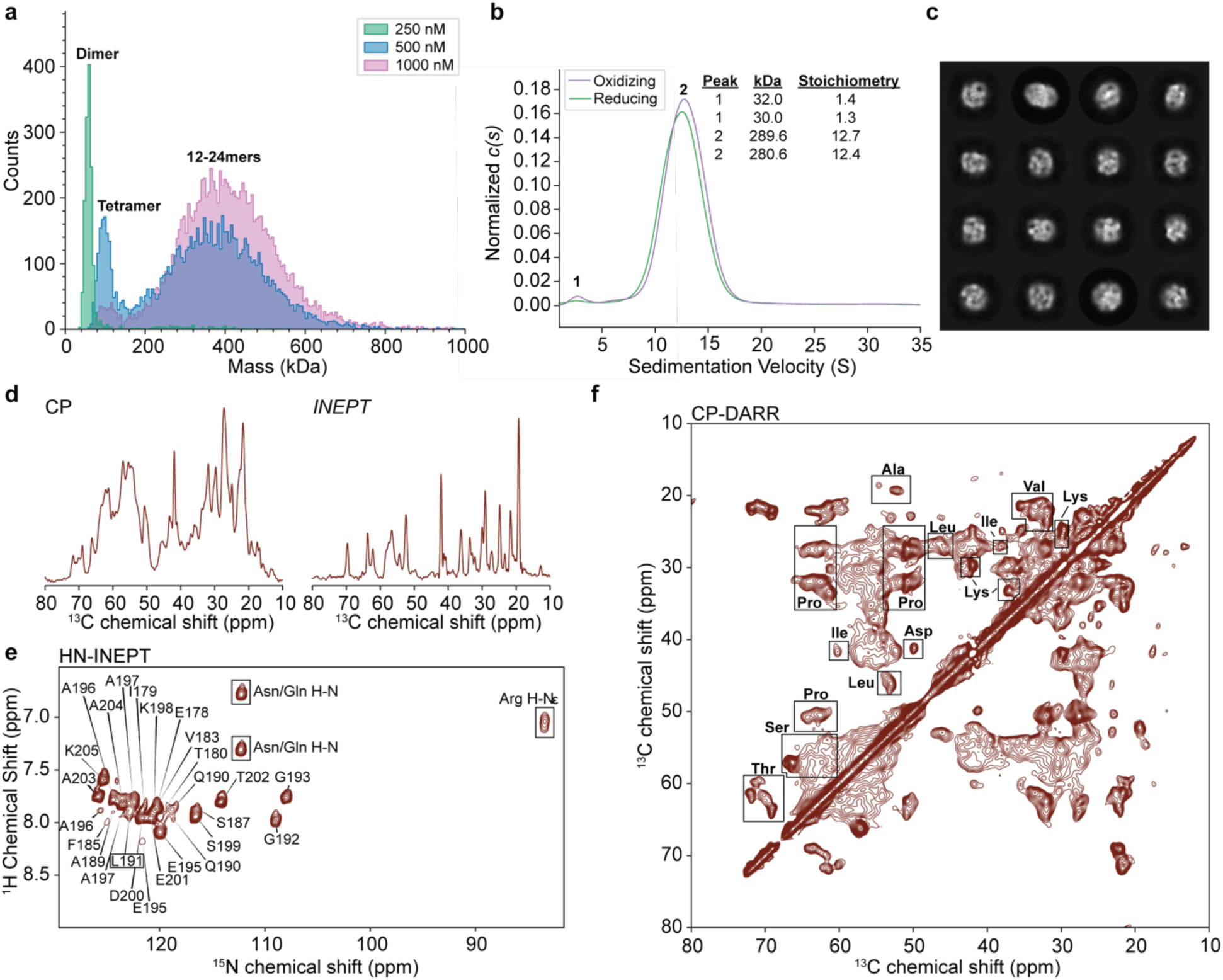
Biophysical and MAS NMR characterization of HSPB1. **(a)** Mass photometry of HSPB1 samples prepared at different concentrations. **(b)** AUC chromatogram of HSPB1 samples (10 μM) prepared under oxidizing and reducing conditions. **(c)** Sample of 2D cryoEM classes of HSPB1 single particles displaying cage-like architectures. **(d)** 1D CP and INEPT spectra of HSPB1 samples. **(e)** 2D HN-INEPT spectrum of HSPB1 overlaid with assignments from Ref. (16). Boxed cross-peaks correspond to side-chain ^1^H-^15^N correlations. **(f)** 2D CP-DARR spectrum of HSPB1. The regions of characteristic amino acid correlations are denoted with boxes. Spectra were recorded on a 750 MHz NMR spectrometer with MAS frequency of 11 kHz.

To gain insight into the three-dimensional architecture of HSPB1 oligomers, we first turned to cryogenic electron microscopy (cryoEM). sHSPs have proven to be a challenging target for EM-based structural characterization, and there are only a few examples of low-resolution structures of sHSP oligomers that are derived from EM experiments (20, 33, 34). Similar to other sHSPs, our HSPB1 oligomers could be visualized and averaged into 2D classes from cryoEM data. These classes appear to be cage-like with symmetric features (**Fig. 1c**). Despite the clear cage-like structures, the secondary structure could not be resolved in 2D classes, likely due to the heterogeneity and flexibility (i.e., bending or stretching moments) of the oligomer architectures. Reconstruction of 2D classes generated volumes that appeared cage-like but did not perform well during refinement, and we chose not to pursue symmetry imposition to avoid the introduction of bias into our final structures. Instead, we decided to focus our efforts on MAS NMR spectroscopy, which has previously been used successfully to extract local structural information from related high molecular weight biological systems that are both dynamic and heterogenous, including oligomers of the sHSP αB-crystallin (18), proteins in biological condensates (35, 36), and amyloid fibrils in cellular lysates (37).

## HSPB1 oligomers contain both rigid and dynamic components

MAS NMR spectroscopy leverages the dipolar couplings present in biomolecular assemblies with long rotational correlation times to report on their structure and dynamics (38). The basic building block of an MAS NMR experiment is the cross-polarization (CP) pulse sequence, which uses a dipolar coupling-based transfer mechanism that is only efficient for rigid components in the sample (39, 40). Solution-type NMR experiments based on the insensitive nuclei enhancement by polarization transfer (INEPT) pulse sequence can be performed on the same sample under MAS conditions to capture dynamic components (41). The combination of these experiments enables the structural and dynamic characterization of complex samples that exhibit motions on multiple time scales (42, 43).

To characterize the local structure and dynamics of HSPB1 in the absence of a client protein, we performed CP and INEPT-based MAS NMR experiments on samples of full-length wild-type HSPB1. **Fig. 1d** presents 1D CP and INEPT spectra demonstrating the presence of both rigid and dynamic components in the sample, while **Fig. 1e** and **Fig. 1f** show the respective 2D INEPT-based ^1^H-^15^N and 2D CP-based ^13^C-^13^C correlations. Of note, the chemical shifts of the correlations in the HN-INEPT experiment match the published assignments of the CTD as determined by solution NMR spectroscopy (**Fig. 1e**) (16). Similar to previous observations (16), this suggests that the CTD in our wild-type oligomeric HSPB1 samples is highly dynamic. On the other hand, the CP-based ^13^C-^13^C spectrum (**Fig. 1f**) is much more complex, presenting a superposition of correlations that are broad and correlations that are relatively well-defined. The spectrum contains signatures that are consistent with the distinct amino acid compositions of the NTD (e.g., enriched in Ala and Pro amino acids) and the ACD (e.g. Lys and Thr amino acids) (**Fig. 2a**), suggesting that both domains are relatively rigid in the context of client-free HSPB1 oligomers. However, the broad features and overlap in the spectrum precluded detailed analysis of each domain, necessitating a different approach as described below.

**Figure 2.**
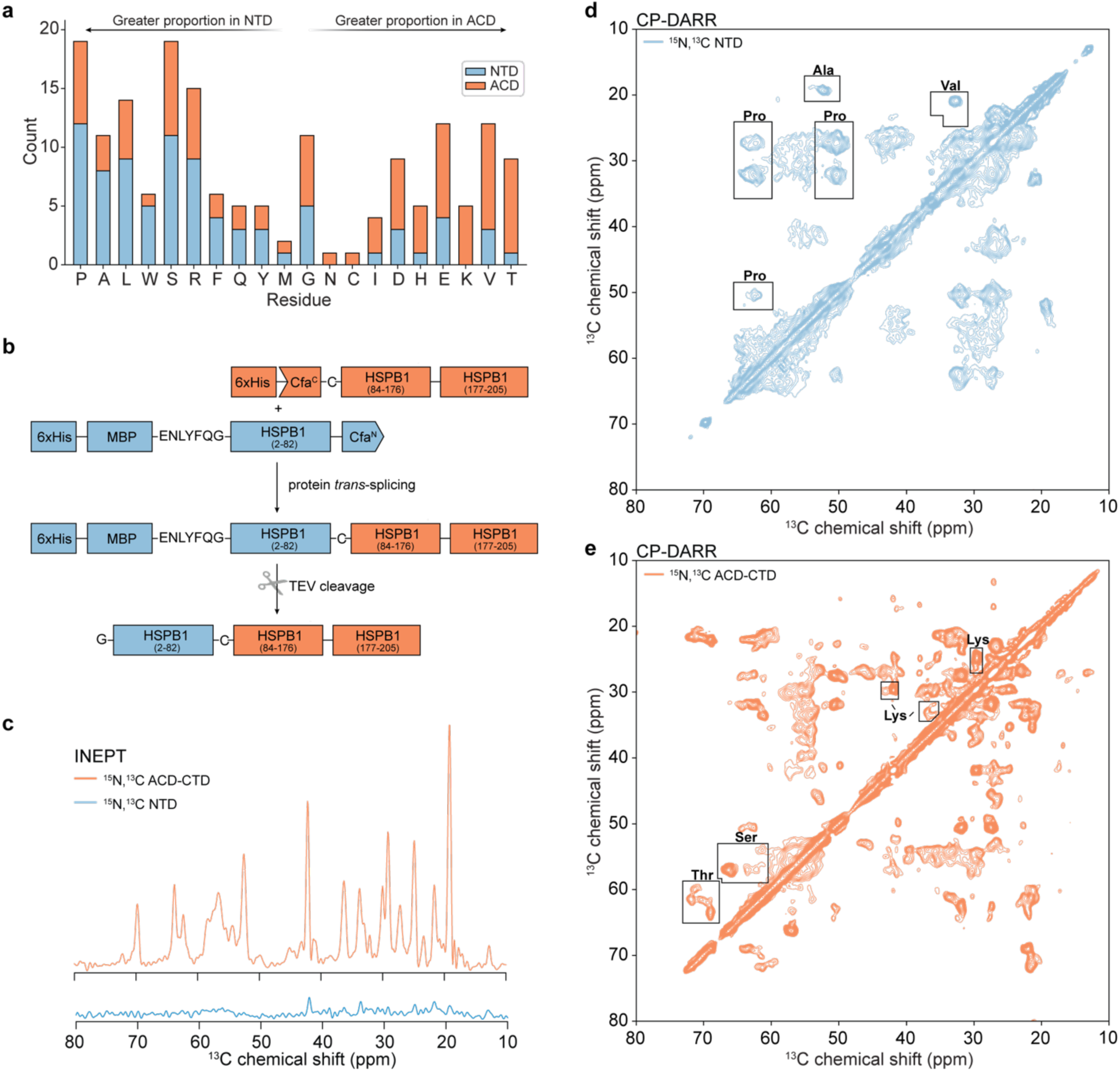
Segmental labeling of HSPB1 facilitates the analysis of MAS NMR spectra. **(a)** Distribution of residues in the NTD and ACD of HSPB1 ordered by relative abundance in each domain. **(b)** Overview of the approach for split-intein mediated segmental labeling of HSPB1. **(c)** 1D INEPT spectra of NTD- and ACD-CTD labeled HSPB1 samples. **(d)** CP-DARR of NTD-labeled HSPB1. Regions of the spectrum containing correlations for residues diagnostic for the NTD are boxed. **(e)** CP-DARR of ACD-labeled HSPB1. Regions of the spectrum containing correlations for residues diagnostic for the ACD are boxed. Spectra were recorded on a 750 MHz NMR spectrometer with MAS frequency of 11 kHz.

## Segmental labeling facilitates unambiguous characterization of HSPB1 domains

To unambiguously characterize the NTD and ACD by MAS NMR, we took advantage of split inteins to prepare a segmentally labeled HSPB1 oligomeric sample (**Fig. 2b**). Split inteins are naturally occurring protein pairs that can carry out polypeptide splicing reactions both *in vitro* and *in vivo* (44). In this process, they build a contiguous protein from two polypeptide segments with minimal changes to the sequence. Inteins are powerful tools for NMR spectroscopy due to their ability to generate segmentally labeled proteins from two recombinant fusion proteins expressed in different media (45).

Here, we chose to use the Cfa_GEP_-engineered split intein system which exhibits rapid kinetics in a range of splicing conditions and only requires a cysteine residue “scar” at the splicing junction of the desired protein (46, 47). Since our goal was to distinguish between the NTD and the ACD contributions in the dipolar-based MAS NMR spectra of HSPB1, we chose to perform splicing at position 83 of the sequence, which required an S83C mutation. We expressed and purified two separate constructs, 6xHis-MBP-HSPB1(1–82)- Cfa^N^ and 6xHis-Cfa^C^-HSPB1(S83C-205), and performed splicing reactions to construct full-length HSPB1 which was either ^13^C,^15^N-labeled on the NTD or the ACD-CTD domains (**Fig. 2b**). The MBP fusion tag was necessary to enhance the otherwise poor solubility of the NTD-Cfa^N^ construct but could be removed in the final step of the purification protocol. This strategy enabled the preparation of 8-10 mg of segmentally labeled full-length HSPB1 for MAS NMR studies (**Figs. S5, S6**). We then proceeded to record INEPT-based and CP-based experiments of the NTD-labeled and the ACD-CTD-labeled samples.

The NTD-labeled sample does not show significant INEPT signals (**Fig. 2c**), but gives robust CP signals, indicating that the NTD is rigid in the oligomeric state. The 2D CP-based ^13^C-^13^C experiment (**Fig. 2d**) shows relatively high sensitivity but also significant broadening, which suggests high degree of heterogeneity in the NTD conformational landscape. Despite the line-broadening, the signals of several amino acid types that are enriched in the NTD can be identified, including Pro, Ala, and Val correlations. Comparison of the chemical shifts of those cross-peaks with chemical shift distributions derived from the Biological Magnetic Resonance Data Bank (BMRB) suggests that the conformations adopted by these residues are consistent with a random coil (**Figs. S7**, **S8**). However, the Ala Cα-Cβ correlations also have a minor component that matches well with the α-helical distribution. Taken together, these results indicate that in the context of HSPB1 oligomers, the NTD domains are rigid but adopt heterogeneous conformations including some α-helical structure.

The spectra of the ACD-CTD labeled sample, on the other hand, have signals consistent with both rigid and dynamic components (**Fig. 2c, e**). The dynamic components are identical to those of the fully labeled HSPB1 sample (**Fig. S9**) and correspond to residues in the CTD domain. The 2D dipolar-based ^13^C-^13^C spectrum (**Fig. 2e**) is relatively well resolved and contains correlations that are characteristic of residues enriched in the ACD domain, including Lys, Thr and Ser. The chemical shifts of these residues are consistent with β-sheet structure (**Fig. S10**). Therefore, in the context of oligomers, the CTD remains disordered and dynamic while the ACD is rigid and structured.

### Fast ^1^H-based MAS NMR spectroscopy captures heterogeneous environments in the ACD

To obtain a more detailed view of the structure and interactions of the rigid components within HSPB1 oligomers, we recorded ^1^H-^15^N and ^1^H-^13^C correlations at fast MAS frequencies of 160 kHz. Under these conditions, the fast MAS spinning significantly attenuates the dipolar couplings between the ^1^H spins, resulting in relatively sharp ^1^H correlations even for rigid and fully protonated systems. Combined with segmental labeling, this allowed us to record dipolar-based ^1^H-^15^N (**Fig. 3a**) and ^1^H-^13^C (**Fig. S11**) spectra of the rigid ACD domain in the context of HSPB1 oligomers. In addition to the greater sensitivity per unit sample enabled by the higher gyromagnetic ratio of the ^1^H spins, these experiments are also valuable reporters on the interactions of backbone and sidechain atoms in the rigid portions of the protein. We chose to focus our fast MAS NMR efforts on the ACD rather than the rigid NTD due to the higher resolution exhibited by this domain at lower MAS frequencies (**Fig. 2d, e**). We also contrast these experiments to the solution-type ^1^H-^15^N experiments of the CTD (**Fig. 1e**), which can be acquired at low MAS or even without spinning due to the fast rotational correlation times exhibited by this domain.

**Figure 3.**
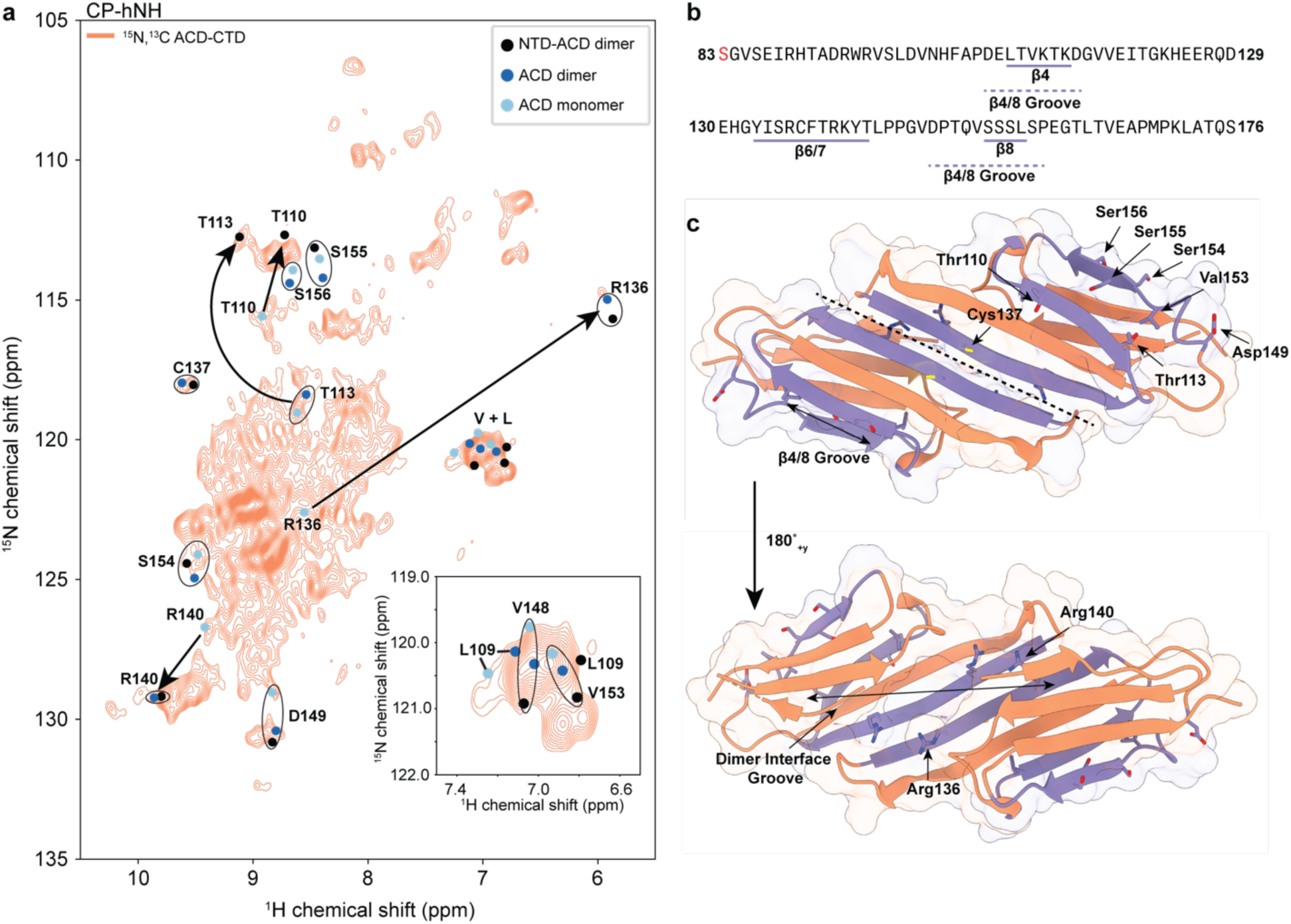
ACD characterization by fast ^1^H-based MAS NMR spectroscopy. **(a)** Dipolar based hNH spectrum of ACD-CTD segmentally labeled HSPB1 oligomers. The solution NMR chemical shifts for diagnostic residues in ACD monomers (Ref. (49)), ACD dimers (Ref.(10)) and NTD-ACD dimers (Ref.(12)) are shown in light blue, dark blue and black, respectively. Significant chemical shift differences are denoted by arrows. The spectrum was recorded on an 800 MHz NMR spectrometer with MAS frequency of 160 kHz. **(b)** Relevant sequence of the ACD domain showing the position of the β4, β6/7 and β8 strands. **(c)** ACD crystal structure (PDB ID: 4MJH, Ref.(11)) depicting the positions of diagnostic residues along the dimer interface and the edge grooves.

Analysis of the dipolar-based ^1^H-^15^N spectrum of the segmentally ^13^C, ^15^N-labeled, fully protonated ACD-CTD domain sample acquired at 800 MHz ^1^H Larmor frequency and 160 kHz MAS (**Fig. 3a**) revealed multiple resolved peaks and a crowded center of the spectrum. While we could not obtain sequential residue assignments at this time, the resolved resonances still provide insight into the ACD chemical environments. To understand these environments, we overlaid the spectrum with ACD assignments obtained by solution NMR spectroscopy (**Fig. S12**). We chose to compare solution NMR assignments from a sample containing ACD monomers only (stabilized by a C137S mutation and low pH) (48, 49), ACD dimers only (10), or NTD-ACD dimers where the NTD contained phosphomimic mutations (S15D, S78D, S82D) to avoid oligomerization (12).

We first sought to confirm that the ACD dimer interface is preserved in the context of HSPB1 oligomers. To this end, we focused on the chemical shifts of several diagnostic residues along the dimer β-sheet interface, namely R136, C137, and R140 (**Fig. 3b,c**). As evident from **Fig. 3a**, these residues exhibit dramatic chemical shift changes from the monomeric to the dimeric form, and the dipolar-based ^1^H-^15^N spectrum clearly shows resolved correlations that are consistent with the presence of a dimer interface. Since these correlations appear in the dipolar-based MAS NMR spectrum, they must be representative of larger oligomer structures, rather than a minor population of small molecular weight dimers that may also be present in the sample. At the same time, due to the position of the monomeric correlations for R136 and R140 (note C137 is mutated to a serine residue in the monomer), we cannot exclude the presence of monomeric-like environments in our samples in addition to the dimer.

Interactions of the NTD and CTD with the edge grooves of the ACD dimer have been implicated in oligomer integrity and chaperone activity (15). In light of these studies, we next asked whether the edge grooves of the ACD domains are occupied in the oligomeric state by analyzing the chemical shifts of several diagnostic groove residues. For example, in previous solution NMR spectra (10, 12, 49), edge groove residues T110 and T113 (**Fig. 3b,c**) display very different chemical shifts in dimer constructs with and without the NTD domain, suggesting that the NTD domain may occupy the edge grooves. Our dipolar-based ^1^H-^15^N spectrum (**Fig. 3a**) shows peaks in both locations, suggesting that both occupied and unoccupied edge grooves are present in HSPB1 oligomers. Other diagnostic residues with unique and well resolved peaks such as D149, V153 and L109 also show chemical shifts consistent with both occupied and empty groove conformations. Finally, the diagnostic groove “bump” residue S155 and its neighbor S156 do not have easily identifiable matches in our spectrum, suggesting that these residues experience different chemical environments compared to their counterparts in the three solution NMR constructs we analyzed (**Fig. S12**).

Taken together, our analysis of the ^1^H-based fast MAS NMR data indicates the presence of intact dimer ACD interfaces and multiple chemical environments for diagnostic edge groove residues, suggesting a complex and heterogenous network of interactions within HSPB1 oligomers.

### The NTD interacts with the ACD grooves and mediates HSPB1 oligomerization

To gain additional insights into the interactions that drive HSPB1 oligomer assembly in the absence of a client, we turned to MD simulations. The literature consensus for the structure of HSPB1 in various states of oligomerization is encoded in AlphaFold2 predictions (**Fig. 4a**) (29, 30). Predictions for the structure of the dimer consistently return high predicted local distance difference test (pLDDT) and predicted aligned error (PAE) scores for the β-sheet-rich ACD and low scores for the CTD, which is consistent with a high degree of order in the ACD and an intrinsically disordered CTD (**Fig. 4b, c)**, similar to what we observe experimentally (50–52). Interestingly, AlphaFold2 also predicts some α-helical secondary structure in the NTD domain, albeit with low pLDDT and PAE scores (**Fig. 4a, b, c**) (53, 54). Predictions for higher order oligomers generate architectures that are based on the HSPB1 dimer. In nearly all cases, oligomer predictions position the NTD towards the center of a cage-like structure (**Fig. 4a**). To further our understanding of the contacts that mediate oligomer integrity, we performed a series of microseconds-long MD simulations with HSPB1 in various states of oligomerization, including monomers, dimers, and dodecamers.

**Figure 4.**
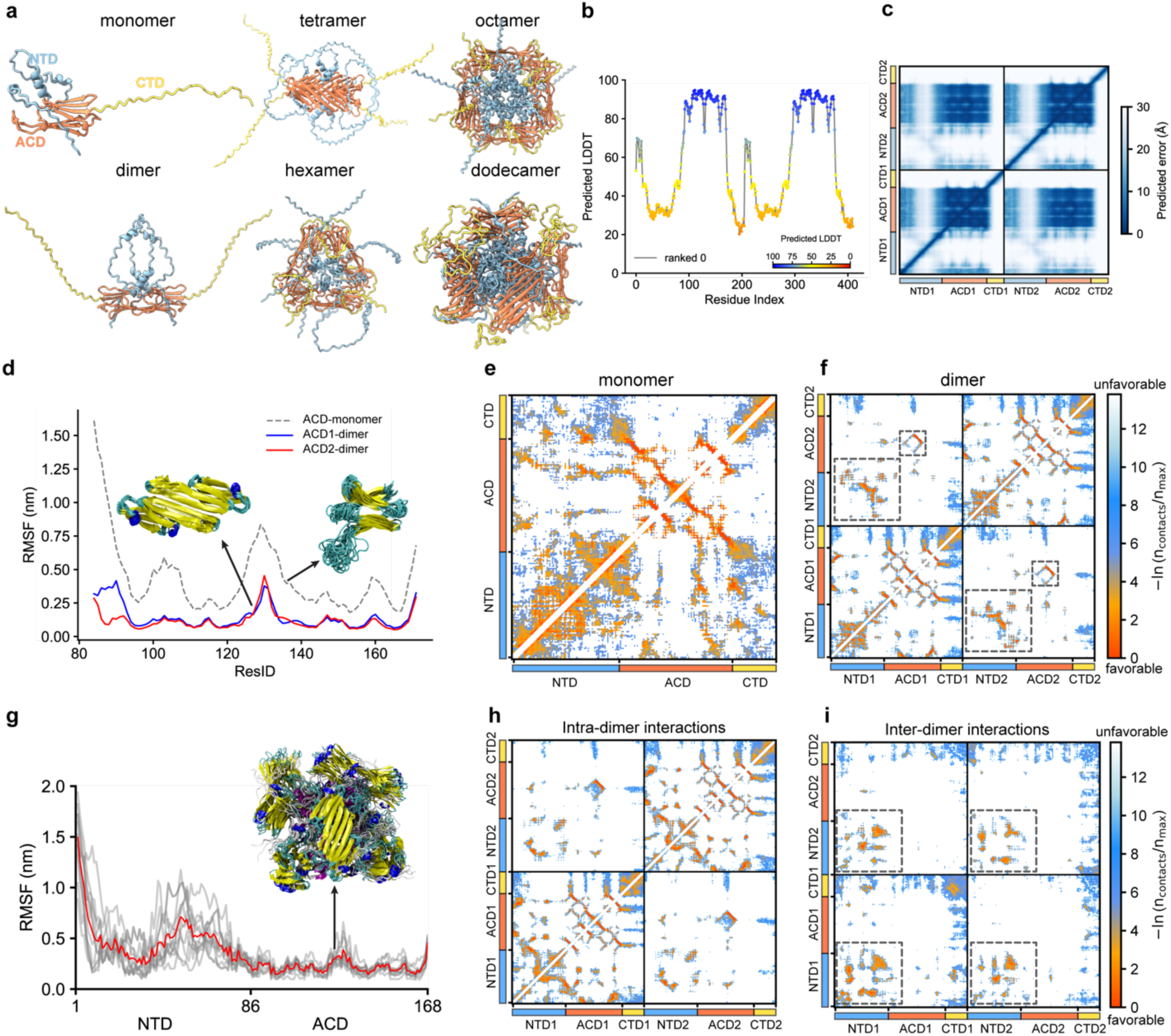
Molecular dynamics simulations reveal the interactions responsible for HSPB1 oligomer assembly. **(a)** Structures of HSPB1 oligomers predicted by Alphafold2. **(b)** pLDDT scores for the HSPB1 dimer prediction. **(c)** PAE scores for HSPB1 dimers with dark blue color indicating lower errors (higher confidence in the predicted distance) and white colors indicating higher errors (lower confidence). **(d)** Root mean-square fluctuation (RMSF) of HSPB1 ACD monomer and dimer forms in all-atom (AA) simulations. Inset shows the conformational ensembles in the AA simulations. The structures are color-coded by secondary structure: purple for α-helix, blue for 3_10_-helix, yellow for β-sheet, cyan for turn/bend, and silver for coil. ACD-1 and ACD-2 refer to each ACD monomer within the dimer form. **(e)** Intramolecular contact map for AA simulations of the monomer. **(f)** Intra-monomer (diagonal quadrants) and inter-monomer (off-diagonal quadrant) contact maps for AA simulations of the dimer. **(g)** RMSF of the NTD and ACD domains in HSPB1 dodecamers obtained from AA simulations. The red line is averaged over the 12 monomers (gray lines) within the dodecamer. **(h, i)** Intra-dimer and inter-dimer contact maps within the dodecamer. The data were averaged from the diagonal and off-diagonal blocks of **Fig. S17a**, respectively. Black boxes highlight favorable interactions.

Previous literature has suggested that HSPB1 monomers are potent chaperones that exhibit local unfolding (48). Consistent with these reports, we found that HSPB1 monomers are relatively unstable in the simulations, with large structural fluctuations occurring in the region comprising β5 and β6+7 (residues 120 – 140) and significant dynamics in the NTD (**Fig. 4d, Movie S1**). The NTD in the monomer adopts transient α- helical secondary structure (**Fig. S13a**) and loosely interacts with large areas of the ACD, including the β4/β8 edge groove (**Fig. 4e, Fig. S14, Movie S1**), whereas the CTD competes for interactions near the β3 strand and the L7/8 loop.

HSPB1 is generally more stable in the dimer state (**Fig. 4d, Movie S2**). Although there is still some motion at the L5/6 loop, the ACD no longer exhibits partial unfolding, and the composite β-sheet formed across the ACD-ACD dimer experiences minimal fluctuations along the 5 μs trajectory (**Fig. 4d**). The NTD adopts a more stable α-helical secondary structure in the region encompassing residues 70 – 80 (**Fig. S13b**). This region is enriched with alanine residues and is adjacent to the insertion NTD sequence that distinguishes HSPB1 from other sHSPs (12). While the NTD makes more specific contacts with the β4/β8 cleft of the ACD, interactions between the NTDs of each monomer are now a prominent feature of the dimer trajectory (**Fig. 4f**). ACD contacts are primarily mediated by the distal (1–18), conserved (25–37) and boundary regions of the NTD (74–86) (**Fig. S15a, c**), while most NTD residues other than the distal region appear to participate in NTD-NTD interactions (**Fig. S16**). Similar to the monomer case, the CTD interacts near the β3 strand and the L7/8 loop, i.e., in the vicinity of the dimer and edge grooves (**Fig. S15b, d**). In addition, intra-monomer CTD-ACD interactions have a much higher probability compared to inter-monomer contacts.

Due to the large size of the system (∼2500 residues), AlphaFold-Multimer (30) (AFM) was unable to predict the full-length HSPB1 dodecamer. For smaller oligomeric structures, we observed that the CTD is disordered and does not participate in the formation of the cage-like structures (**Fig. 4a**). Therefore, to reduce the system size, we ran AFM predictions on the truncated HSPB1 dodecamer (HSPB1-ΔCTD) with and without templates (known protein structures used as references to aid in the prediction as described in the Materials and Methods). Models predicted without templates formed symmetric and complete cages, using dimer structures as the fundamental units (**Movie S3**). In contrast, models predicted with templates (aligned from PDB 6dv5(55)) predominantly adopted structures resembling half of a 24mer, with monomers serving as the fundamental units (**Movie S4**). In the highest ranked structures predicted without a template, the relative orientation of each HSPB1 dimer within the dodecamer was different. Nevertheless, all dodecamer structure predictions exhibited cage-like architectures in which the NTD was sequestered in the cage, the ACD dimer formed the cage, while the side chains of the ACD lysine residues faced towards the solvent.

To explore the role of HSPB1’s domains in the oligomer state, we used Modeller(56) to add the CTD to the top-ranked model of HSPB1-ΔCTD (predicted without templates) and performed all-atom simulations for 5 μs (**Movie S5**). As was the case with the dimer, many of the core features of the HSPB1 dimer persist along the trajectories of dodecamer configurations – the ACD remains stable and intact, the NTD makes specific contacts with the β4/β8 grooves of the ACD within each dimer building block, while the CTD is disordered and dynamic (**Fig. 4g, h, I**). However, the NTD’s role as a mediator of high-order oligomerization becomes more apparent in the dodecamer simulations. The interactions within and between dimers, mediated by the aromatic (residues 15 – 20) and tryptophan-rich (residues 38 – 44) regions, as well as the α-helical region (residues 70 – 80) in the NTD, contribute to the stability of the dodecamer core (**Fig. 4h, i** and **S17**, **S18**). While the NTD-NTD interaction patterns are somewhat different within the dimer building block and between dimers, in all cases they exclude the distal region of the NTD (**Fig. S17c, d**). In our model, the CTD does not appear to interact extensively with other domains in the dodecamer and remains dynamic throughout the simulations (**Fig. 4h, i**).

### HSPB1 oligomer size decreases in the presence of a client protein

Our next goal was to characterize the wild-type HSPB1 oligomers in the presence of a client protein using mass photometry experiments and MD simulations. As a model client protein, we chose FUS LC whose interactions with HSPB1 dimers have previously been characterized by solution NMR spectroscopy (13). Samples of 0.5 μM HSPB1 alone show mass photometry distributions consistent with tetramers and higher order oligomers (**Fig. 5a**). On the other hand, FUS LC monomers are ∼17 kDa and hence below the detectable range for mass photometry. Nevertheless, FUS LC only samples appear to have distributions in the 100 – 152 kDa range (**Fig. S19**), suggesting protein oligomerization in the absence of HSPB1 and at protein concentrations that are below the saturation concentration for condensate formation, similar to previous observations (57). In mixed samples containing both HSPB1 and FUS LC, we observed two major trends. First, as the concentration of FUS LC increased, the lower molecular weight peak initially centered around 85 kDa increased in intensity and moved to a higher molecular weight distribution centered around 160 kDa. Based on the molecular weight distribution, this peak could contain a mixture of FUS LC oligomers, lower molecular weight HSPB1 oligomers, and interaction complexes between the two proteins. Second, as the concentration of FUS LC increased, the intensity of the high molecular weight oligomeric peak (centered around 440 kDa) decreased, suggesting that the relative proportion of higher-order HSPB1 oligomers was reduced as more client protein was available for interactions. This observation implies that HSPB1 oligomers reorganize into lower molecular weight species in the presence of a client protein.

**Figure 5.**
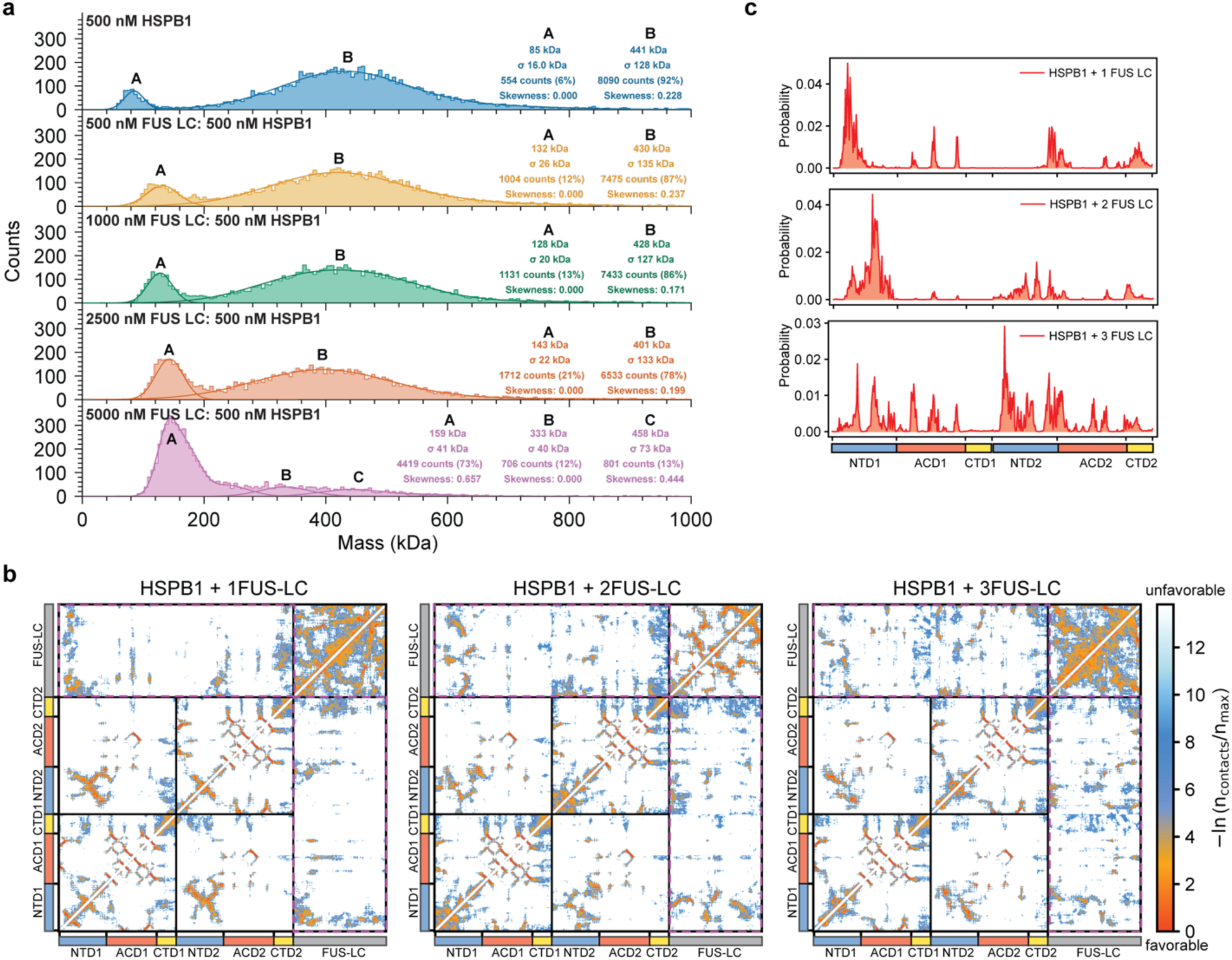
Interactions of HSPB1 with FUS LC under dilute conditions. **(a)** Mass photometry gaussian-fit analysis of HSPB1 (500 nM) in the presence of increasing concentrations of FUS LC (0-5000 nM). Measurements were performed in a 150 mM NaCl and 50 mM sodium phosphate (pH=7.2-7.4) buffer. **(b)** AA simulations contact maps for HSPB1 dimers in the presence of one, two and three FUS LC monomers. **(c)** Summed contact probabilities for each residue in the HSPB1 dimers (rectangle box) interacting with one, two or three FUS LC monomers.

To explore the interactions between HSPB1 and the client protein, we extended our all-atom MD simulations of the HSPB1 dimer to include one, two, or three FUS LC monomers (**Movie S6-8**). To facilitate interactions, FUS LC chains were positioned at different locations within the simulation box, maintaining a minimum distance of 8 Å from HSPB1. As shown in **Fig. 5b**, FUS LC engages both the NTD and the ACD of the HSPB1 dimer, with occasional contacts to the CTD. The interactions between FUS LC and ACD become increasingly prominent as more FUS monomers are included in the simulation (**Fig. 5c, S20a**). FUS LC-NTD interactions are clustered in two regions across residues 5-21 (distal and aromatic regions) and 28-38 (conserved region) in the NTD, and two regions across residues 25-36 and 47-60 in FUS LC (**Fig. S20b, c**). Interestingly, the interaction regions on the FUS LC sequence are not part of the β-strands that form the core of FUS LC fibrils, generally lying either just outside the fibril-forming region or existing as loops in between RAC domains or in between β-sheets in the case of Type I and Type III fibril conformations (58–60). Similar to tau, FUS LC interacts with the β4-β8 groove in the ACD in our simulations (15, 22). It also engages with the adjacent β5 and β6+7 ACD β-sheets, mediated by tyrosine residues in FUS LC (**Fig. S20c, d**), a result that is consistent with experiments in the literature that have been performed with FUS LC and an ACD construct of HSPB1 (13).

In summary, in the presence of the client protein FUS LC under dilute conditions, the average size of the HSPB1 oligomers decreases, favoring lower molecular weight heterogenous assemblies that are ∼160 kDa in size. This is consistent with literature suggesting that lower molecular weight HSPB1 species are more active chaperones (3, 7). Similar to the published literature, our simulations of dimers and FUS LC monomers suggest interactions through both the NTD and ACD domains in HSPB1 (13).

### The NTD is essential for HSPB1 dimer partitioning into FUS LC condensates

We next wondered how HSPB1 oligomers engage clients in the context of biological condensates. Previous literature has suggested that wild-type HSPB1 can disrupt FUS LC droplet formation (13). Consistent with these findings, we observed similar behavior in our experiments. At a 5:1 ratio of FUS LC to HSPB1, the two proteins co-localize into droplets, while increasing amounts of HSPB1 disperse the droplets (**Fig. S21**). We also prepared droplets containing unlabeled FUS LC and segmentally isotopically labeled HSPB1 and performed ^13^C T1ρ relaxation MAS NMR experiments. The T1ρ relaxation parameter, which is a useful measure of μs-ms dynamics, decreased for both the NTD- and ACD-labeled HSPB1 samples compared to samples prepared with only chaperone (**Fig. S22**), This suggests that both the NTD and ACD experience changes in dynamics in the condensed phase. These changes may arise from an altered oligomerization state of the chaperone, interactions of the two domains with the client protein, and the different viscosity of the droplet environment.

To capture the potential interactions of HSPB1 in FUS LC condensates, we performed coarse-grained (CG) co-existence simulations using the HPS-Urry model that has previously been shown to faithfully recapitulate LLPS *in silico* (32, 61, 62). In the simulations, we used a mole fraction of FUS LC to HSPB1 of 2:1 and kept the ACD domains rigid with respect to each other in the dimer configuration, while the NTD and CTD remained flexible (see Materials and Methods). We first compared the ability of HSPB1 monomers, dimers, hexamers and dodecamers to partition into FUS LC droplets and observed that dodecamers lead to higher concentrations of FUS LC in the dilute phase (**Fig. S23**), suggesting that they attenuate phase separation. In contrast, HSPB1 dimers fully partition into FUS LC condensates, without affecting the phase separation propensity of the client protein (**Fig. 6a, Movie S9**). This is evident by the same dilute concentration of FUS LC in condensates whether in the presence or absence of HSPB1 (**Fig. 6a**). These observations are consistent with previous experiments that have shown the ability of dodecamers to disrupt FUS LC droplets while mutation-stabilized HSPB1 dimers partition into FUS LC droplets at much higher concentrations compared to the oligomers (13). To characterize the molecular interactions involved in co-partitioning, we computed the 1D and 2D time-averaged intermolecular contact map between FUS LC and HSPB1 dimers as a function of residue number (**Fig. 6b**). The contacts in FUS LC are relatively evenly distributed, with intermittent peaks corresponding to tyrosine residues throughout its sequence. In contrast, the contacts in HSPB1 are predominantly contributed by the NTD, with additional contributions from the CTD. Some regions in the ACD show diminished contacts due to the occlusion caused by the enforced ACD fold (**Fig 6b**).

**Figure 6.**
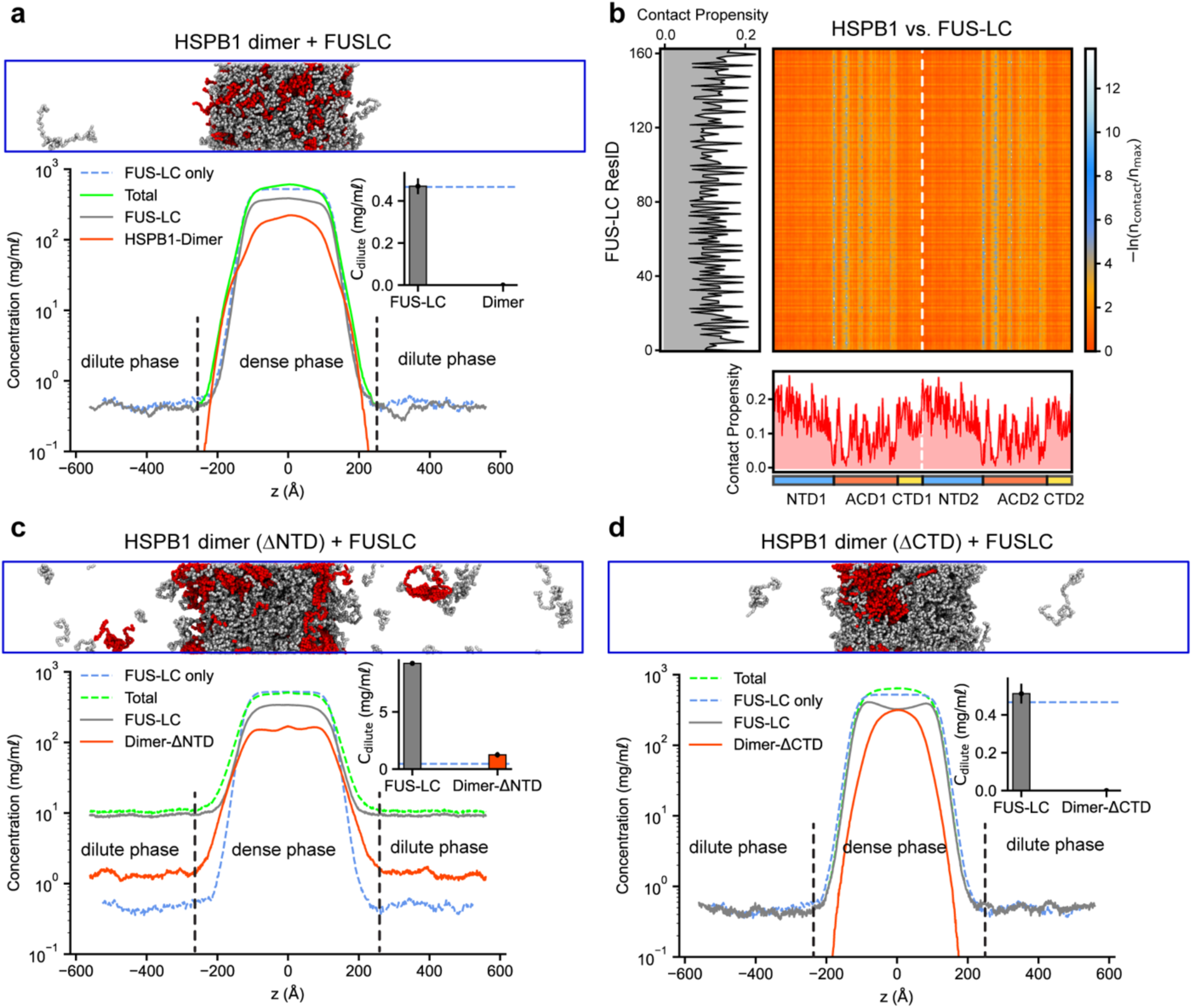
CG phase coexistence simulations suggest an essential role for the NTD in HSPB1 dimer partitioning into FUS LC condensates. **(a)** Density profiles, dilute concentrations (inset), and a snapshot of FUS LC and HSPB1 in CG phase coexistence simulations. **(b)** Intermolecular contact map between HSPB1 and FUS LC within the condensed phase. Preferential interactions are shown in red. The 1D contact maps at the bottom and on the left show the average contact propensity per frame per residue for the HSPB1 dimer and FUS LC, respectively. **(c, d)** Density profiles, dilute concentrations (inset), and snapshots of HSPB1 and FUS LC upon removal of NTD and CTD domains, respectively. The dashed blue lines in the insets denote the dilute protein concentration from a simulation that includes only FUS LC.

We next explored the effects of the NTD and CTD on the co-partitioning properties of HSPB1 dimers. Simulations of FUS LC and an HSPB1-ΔNTD construct reveal an increase in the dilute phase concentration of HSPB1 (**Fig. 6c**). In addition, the dilute concentration of FUS LC increases by more than an order of magnitude, while the overall condensate density is reduced (**Fig. 6c, Movie S10**). Removal of the CTD of HSPB1, in contrast, does not have a significant effect on the formation of the condensate (**Fig. 6d, Movie S10**). In agreement with our observations in all-atom simulations of FUS LC and HSPB1 and previous experimental results(13), this suggests that the NTD plays a significant role in engaging FUS LC clients and further indicates that the NTD is responsible for the partitioning of HSPB1 into FUS LC droplets and in promoting the integrity of the FUS LC-HSPB1 co-condensates.

## Discussion

In this study, we leverage split-intein mediated segmental labeling, ^13^C, ^15^N and ^1^H based MAS NMR spectroscopy, and MD simulations, to characterize the structure, dynamics and interactions of the HSPB1 chaperone, both in the presence and absence of a client protein. The combination of these approaches has allowed us to shed light on the elusive wild-type state of the protein, which forms polydisperse cage-like oligomers that contain both rigid and dynamic components and has been difficult to characterize experimentally at high resolution. Our results show that within the HSPB1 oligomers, the CTD tails are dynamic and disordered, the NTDs are rigid and highly heterogenous, while the ACDs exist as rigid β-sheet-rich dimers. Our results also indicate that wild-type HSPB1 oligomers do not partition efficiently into client condensates and that the NTD domain is required for partitioning. In addition, higher order HSPB1 oligomers appear to restructure into smaller oligomeric species in the presence of a client protein. Below, we discuss these observations in more detail.

Regarding the CTD, our observations are consistent with the idea that this domain serves as a solubility tag for HSPB1 oligomers (12). In our MAS NMR experiments, we observe the CTD only in solution-type experiments, suggesting that this domain is highly dynamic. ^1^H-^15^N and ^1^H-^13^C INEPT spectra indicate that it adopts a random coil conformation, similar to those observed by others in the context of oligomers or in mutation-stabilized dimers (14, 16). Our simulations indicate that except for some CTD-CTD contacts, this domain does not participate in extensive interactions in oligomers, however, some ACD-CTD interactions can be detected in dimers and monomers. Although the ACD-CTD interaction patterns in these cases do not match exactly the specific interaction between the IXI motif in the CTD and Ser155 in the edge groove detected by solution NMR studies (14), our simulations indicate that the CTD explores the neighboring loop regions. We also note that we cannot exclude the presence of IXI- Ser155 interactions in our MAS NMR samples, as they may not be detectable due to low populations or intermediate timescale dynamics that would make such species invisible to both solution and solid-state type experiments.

The combination of segmental labeling and MAS NMR spectroscopy allowed us, for the first time, to observe the rigidity and heterogeneity of the NTD in the absence of any mutations or modifications. While the chemical shifts for most peaks appear to indicate random coil conformations, there is strong indication that alanine residues also adopt α-helical structures. Alanine residues are particularly enriched in the NTD insertion region (57–70) that is unique to HSPB1, while our simulations detect strong α-helical propensity for residues 70-80 (boundary region) which also includes two alanine residues. Considering the heterogeneity in our NTD spectra, it is likely that both the α-helical and random coil conformations exist at the same time. While some residues in the 70-80 region may participate in ACD contacts in the context of dimers, the insertion and boundary regions feature prominently in inter-dimer NTD-NTD interactions in the oligomer simulations, suggesting that the α-helical conformation may be important for oligomer formation and the sequestering of the NTD tails. Previous work has also suggested that other NTD regions may experience transient α-helical conformations (e.g. the Trp-rich region) (12), however, we currently lack the experimental resolution to detect those. In addition, α-helical NTD regions have also been detected in the closely related HSPB5 chaperone where they were modeled as structured components within the oligomers (17, 19).

Our MD simulations indicate that NTD contacts are important in the interaction maps of HSPB1 monomers, dimers and 12-mers. While ACD-NTD contacts persist throughout all interaction maps, the 12-mer maps are dominated by NTD-NTD contacts, reinforcing the idea that these interactions play an important structural role in oligomer assembly. In all template-free AlphaFold2 models of HSPB1 oligomers, the NTDs are sequestered in the interior of the oligomers, creating cage-like structures that are consistent with our cryoEM images. In dimers, the NTD uses the distal, aromatic, conserved and boundary regions to engage with the ACD, conclusions that match well with experimental data from solution NMR of mutation-stabilized dimers (12). On the other hand, all NTD segments except for the distal region can engage in NTD-NTD interactions, also consistent with the conclusions from the solution NMR study (12). In oligomers, the NTD-NTD interactions become the dominant feature, although the distal region continues to be excluded from the interaction maps. These observations suggest that in dimers and smaller oligomers, the NTD may explore the ACD surface more compared to the oligomeric state, where NTD-NTD contacts dominate perhaps as a means to sequester the NTD into the oligomer interior.

We also took advantage of recent developments in fast MAS NMR spectroscopy to record the first ^1^H-^15^N and ^1^H-^13^C spectra of the ACD domains within wild-type HSPB1 oligomers. Despite the inherent heterogeneity of the sample and the peak overlap, the spectra agree quite well with solution NMR spectra of NTD truncation or mutation-stabilized HSPB1 dimers, suggesting that the β-sheet-rich ACD dimer is the main building block of the oligomers. More importantly, however, the data indicate at least two (but possibly more) discrete chemical environments for the edge grooves suggesting that there are empty and occupied groves. Our simulations indicate that there are differences in the interaction patterns of the ACD with the CTD and the NTD domains within the monomers of a dimer, between the monomers of dimers, and between dimers within the oligomer context, highlighting potential sources for the experimentally observed heterogeneity. Nevertheless, whether they are with the CTD or the NTD, the ACD interactions appear centered on the edge or dimer grooves and their vicinity, reinforcing the important role these structural features play in HSPB1 oligomer assembly.

Wild-type HSPB1 appears to undergo a dramatic structural transition in the presence of a model client protein, FUS LC. As shown by mass photometry, the average size of the HSPB1 oligomers decreases, while NMR relaxation measurements indicate enhanced dynamics in the NTD and ACD domains. Taken together, these observations are consistent with a mechanism where the larger oligomers rearrange into smaller, more dynamic units. Our MD simulations indicate that FUS LC can interact with all HSPB1 domains, although NTD contacts dominate under both dilute and condensed phase conditions. These results align well with experimental observations that suggest client proteins such as tau and FUS LC can interact with both the ACD and NTD domains in HSPB1 dimers, however, only the NTD contacts are productive and prevent aggregation (13, 15).

Consistent with previous literature (13), our fluorescence microscopy data indicates that increasing concentrations of HSPB1 disrupt FUS LC droplets. Our simulations also indicate that dodecamers are less efficient at partitioning into condensates compared to dimers and so are dimers that lack the NTD domain. Less partitioning, however, is coupled to higher efficiency of condensate disruption. In our dodecamer simulations, the NTD is sequestered in the center of the cage-like HSPB1 oligomer, and therefore inaccessible to FUS LC, an effect that is mimicked by the HSPB1- ΔNTD construct. Therefore, exposed NTD tails are required for interactions with the client protein and partitioning into condensates. On the other hand, missing or sequestered NTD tails not only result in lack of partitioning but are also more efficient in disrupting condensates. These conclusions align well with experimental data that shows that wild-type HSPB1 disrupts FUS LC condensates at lower ratios, while mutation or phosphorylation stabilized dimers partition into droplets at much higher ratios (13).

Our results build upon published observations that suggest a leading role for the NTD domain in the assembly of the HSPB1 oligomers and their interactions with client proteins (13, 15). In the absence of a client protein, HSPB1 exists in heterogenous cage- like oligomeric assemblies where the NTD domains are potentially sequestered within the cage. We propose that NTD-NTD contacts within the cage are mediated through partially helical segments. The oligomeric form is built of ACD dimers and reinforced by NTD-ACD interactions, while solubility is imparted by the CTD domains. In cells, these assemblies may function as NTD “storage” units or may play a role in the regulation of biological condensates. In the presence of client proteins, HSPB1 oligomers rearrange into smaller units that likely expose the NTD domains. These units can interact with clients more efficiently and are more potent chaperones. In cells, the transition from higher order assemblies to smaller oligomers is likely controlled by NTD phosphorylation, which favors the dimeric form of the chaperone (63, 64).

In summary, our combined spectroscopic, chemical biology, and computational approach has allowed us to visualize the heterogeneous HSPB1 oligomeric landscape at the molecular level, and to being unraveling the complex dynamics and interaction networks of the chaperone and its clients in the LLPS state. We believe that ^1^H-based fast MAS NMR spectroscopy, combined with segmental labeling and MD simulations, will play an essential role in the further characterization of HSPB1 and its clients, as well as many other heterogenous biological systems that contain regions of order, disorder, and “quasi-order”.

## Materials and Methods

Detailed information regarding the materials and methods is presented in the supporting information file. This includes detailed protocols for the AlphaFold predictions, AA MD simulations, CG MD simulations, cloning and expression of HSPB1-related constructs, purification of HSPB1 constructs, mass photometry, analytical ultracentrifugation, negative staining and transmission electron microscopy, purification and splicing of HSPB1 intein constructs, expression of FUS LC, purification of FUS LC, insulin aggregation assay, fluorescent labeling and microscopy of the proteins, MAS NMR experiments and data analysis, cryoEM microscopy sample preparation and data collection, and cryoEM microscopy image processing.

## Acknowledgements

This work was supported by NIH grants R35GM138382 (GTD), R35GM153388 (JM), R35GM138206 (MH), and T32GM139795 (APP). We are grateful for the computational resources provided by Texas A&M High Performance Research Computing (HPRC); the NMR support provided by Dr. Mounesha Garaga Nagendrachar; the mass photometry support provided by Prof. Andres Leschziner; the cryoEM advice and instrumentation provided by Dr. Mariusz Matyszewski, Robert Ashley, and the cryoEM facility at UCSD; and the support provided by Brendan Dennis, Kevin Smith, and the UCSD Physics Computing Facility. The work presented here used facilities supported by NIH grants P30NS047101 (UCSD School of Medicine Microscopy Core) and P30 CA030199 (Sanford Burnham Prebys Protein Analysis Core).

