## Supplementary material for "Capturing the Conformational Heterogeneity of HSPB1 Chaperone Oligomers at Atomic Resolution": Methods and Supporting Figures

### Materials and Methods

#### AlphaFold predictions

The predicted structure of the HSBP1 monomer was obtained from the full-length model available in the AlphaFold protein structure database, hosted by the European Bioinformatics Institute (EBI). https://alphafold.ebi.ac.uk/entry/P04792

AFM v2.3.0 was employed to generate the structures of the HSPB1 dimer, tetramer, hexamer, and octamer (**Fig. 4a**). The AFM computations were executed on a single NVIDIA DGX-A100 GPU and 24 CPUs with 180 GB of memory. Sequence searches were conducted using the --db_preset=full_dbs flag, incorporating the following databases: Uniclust30 (1) (version UniRef30_2021_03), UniRef90 (2) (version 04/24/2022), BDF (3), MGnify cluster (mgy_clusters_2022_05), and the PDB, mmCIF, and SEQRES databases as of January 1, 2022. The quality of the structural models was evaluated using both the mean per-residue predicted Local Distance Difference Test (pLDDT) score and the Predicted Aligned Error (PAE), which indicate local and overall structural accuracy (4, 5), respectively. Models were ranked based on the ranking_confidence score provided by AlphaFold. This score is a weighted average of the interface predicted template modeling score (ipTM) and the overall predicted template modeling score (pTM), calculated as: ranking_confidence = 0.8ipTM + 0.2pTM (6). While pTM measures errors within individual protein chains, ipTM evaluates the accuracy of the predicted interactions between chains. The top-ranked models shown in **Fig. 4a** were colored by pLDDT confidence scores using ChimeraX (7).

Due to an out-of-memory error, AFM was unable to predict the structure of the full-length HSPB1 dodecamer or its truncated dodecamer form (HSPB1-ΔCTD). However, ColabFold (1) version 1.5.2 successfully predicted the structure of the truncated HSPB1 dodecamer. ColabFold was downloaded from GitHub (https://github.com/YoshitakaMo/localcolabfold) along with the environmental databases (https://colabfold.mmseqs.com) and installed on a local computer. ColabFold (alphafold2_multimer_v3) was run locally using 48 recycles, MMseq2 for multiple sequence alignments, and refinement with Amber. Five models were generated and ranked according to their ipTM scores. Full-length HSPB1 dodecamer models were constructed by connecting the disordered C-terminal regions to the truncated models using MODELLER (8).

#### All-atom molecular dynamic simulations

The monomer and top-ranked dimer and dodecamer HSPB1 structures were prepared using Gromacs 2021.6 (9). System topologies were modeled using the Amber03ws force field (10), available at https://bitbucket.org/jeetain/all-atom_ff_refinements. Octahedral boxes with lengths of 11.5 nm, 15 nm, and 17.5 nm were chosen to provide at least 1.5 nm of water molecules around the monomer, dimer, and dodecamer, respectively. In the simulations of the HSPB1 dimer with FUS LC chains, a 17 nm octahedral box was used. The FUS LC chains were inserted at various locations within the box, maintaining a minimum distance of 8 Å from HSPB1 to facilitate interactions between the two components. The system's energy was initially minimized in vacuum, followed by another energy minimization after solvation with TIP4P/2005 water molecules (11), using the steepest descent algorithm. To replicate physiological salt concentration (150 mM), $Na^+$ and $Cl^-$ ions were added along with additional counter ions to achieve electrical neutrality. Improved salt parameters from Lou and Roux (12) were employed for all simulations. The system was first equilibrated for 100 ps in a canonical ensemble (NVT) using a Nose-Hoover thermostat (13) with a coupling constant of 1.0 ps at 300 K, followed by another 100 ps equilibration in an isothermal-isobaric ensemble (NPT) using a Berendsen barostat (14) with an isotropic coupling constant of 5.0 ps to achieve a pressure of 1 bar.

Production simulations were conducted in Amber22. The Gromacs files were converted to Amber inputs using "gromber" in Parmed (15), and hydrogen mass repartitioning to 1.5 amu was performed during the conversion to facilitate a timestep of 4 fs for production runs. After the conversion, a minimization run was performed using the steepest descent and conjugate gradient algorithms, with protein positions restrained. The energy-minimized structures were then heated for 5 ns, with the temperature linearly increasing from 100 K to 300 K during the first half and maintained at 300 K during the second half. Subsequently, two NVT simulations at 300K were conducted: a 5 ns equilibration with restrained protein backbone and an additional 5 ns equilibration with all restraints released. The systems were further equilibrated for 10 ns with a 4 fs timestep in NPT using a Monte Carlo barostat (16) with an isotropic coupling constant of 1.0 ps at a pressure of 1 bar. Temperature control was achieved using Langevin dynamics with a friction coefficient of 1.0 $ps^{-1}$, and bonds involving hydrogen atoms were constrained using the SHAKE algorithm (17). A non-bonded cutoff of 0.9 nm was applied for short-range interactions, while long-range electrostatic interactions were treated using the Particle Mesh Ewald (PME) method (18, 19) Finally, the systems underwent a 5 μs NPT production run with the same parameters as used during equilibration. Distance-based pairwise contact maps were calculated using the MDAnalysis (20) package version 2.5.0, with two residues considered to form a van der Waals contact if at least one atom from one residue was within 6 Å of an atom from the other residue. This distance cutoff, as extensively validated in prior studies (21-23), effectively captures various interaction modes, such as van der Waals interactions, hydrogen bonds, and salt bridges.

**Coarse-grained molecular dynamics simulations**

The coarse-grained (CG) coexistence simulations of HSPB1 and FUS LC were conducted using the HOOMD-Blue 2.9.7 software package (24), following the protocol detailed in our previous studies (23, 25-27). These simulations were run for 5 μs in an NVT ensemble with a Langevin thermostat, employing a friction factor $\gamma = m_{AA}/\rho$, where $m_{AA}$ represents the mass of each amino acid bead and $\rho$ is the damping factor, set to 1000 ps. The timestep was 10 fs. The mole fraction of FUS LC to HSPB1 dimer (or its truncations) was maintained at 2:1, consistent with our experimental setup. In these simulations, FUS LC was modeled as flexible, while the ACD domains of the HSPB1 dimer were constrained as a single rigid body using the hoomd.md.constrain.rigid function (28), allowing the rest of the chain to remain flexible. This ensured that the ACD domains stayed fixed relative to each other in a dimer configuration, while the folded domains were free to move (translate and rotate) and interact with other folded, disordered segments and FUS LC chains during the simulation. The CG coexistence simulations of FUS LC with HSPB1 dimer (and its truncations) and HSPB1 dodecamer were initiated in box dimensions of 180×180×1260 $Å^3$ and 200×200×1400 $Å^3$, respectively. The number of chains of FUS LC and HSPB1 was chosen to give similar protein concentrations in these simulations. The simulation box size and the number of protein chains were selected to account for the potential impact of finite-size effects as done in our previous work (23). All CG simulations were performed at a salt concentration of 100 mM, and a temperature of 300 K. For density profile and contact map calculations, the first 1 μs of the trajectory was excluded as an equilibration period. In the contact analyses, two residues were considered to be in contact if the distance between them was less than 1.5 times the arithmetic mean of their Van der Waals (vdW) radii. Snapshots of the simulations were visualized using VMD (29).

**Cloning and expression of HSPB1-related constructs**

The HSPB1 sequence was sourced from a template plasmid encoding the 6xHis-TEV-HSPB1 construct, a gift from Prof. Don Cleveland, from which the 6xHis- Tobacco Etch Virus (TEV) sequence was removed using an NEBuilder HiFi DNA Assembly Cloning Kit (New England Biolabs, Ipswitch, MA) and custom primers (Integrated DNA Technologies, Inc., Coralville, IA). HSPB1 intein constructs were prepared using HIFI Assembly (New England Biolabs, Ipswich,

MA). The $Cfa_{GEP}$-engineered split intein system was used (30). The resulting intein plasmids in this study encode 6xHis-MBP-TEV-HSPB1[NTD]-$Cfa^N$-6xHis or 6xHis-$Cfa^C$-HSPB1[S83C-ACD-CTD] constructs. In addition, a full-length construct for fluorescent labeling encoding the amino acids GSKCK was cloned into the terminal end of the CTD using an NEBuilder HiFi DNA Assembly Cloning Kit (New England Biolabs, Ipswitch, MA). The resulting plasmid contained the HSPB1-GSKCK construct.

*Full-length HSPB1:*
MTERRVPFSLLRGPSWDPFRDWYPHSRLFDQAFGLPRLPEEWSQWLGGSSWPGYVRPLPPA
AIESPAVAAPAYSRALSRQLSSGVSEIRHTADRWRVSLDVNHFAPDELTVKTKDGVVEITGKHEE
RQDEHGYISRCFTRKYTLPPGVDPTQVSSSLSPEGTLTVEAPMPKLATQSNEITIPVTFESRAQL
GGPEAAKSDETAAK

*6xHis-MBP-TEV-HSPB1[NTD]-$Cfa^N$-6xHis:*
MGSDKIHHHHHHSSGTKIEEGKLVIWINGDKGYNGLAEVGKKFEKDTGIKVTVEHPDKLEEKFP
QVAATGDGPDIIFWAHDRFGGYAQSGLLAEITPDKAFQDKLYPFTWDAVRYNGKLIAYPIAVEAL
SLIYNKDLLPNPPKTWEEIPALDKELKAKGKSALMFNLQEPYFTWPLIAADGGYAFKYENGKYDI
KDVGVDNAGAKAGLTFLVDLIKNKHMNADTDYSIAEAAFNKGETAMTINGPWAWSNIDTSKVNY
GVTVLPTFKGQPSKPFVGVLSAGINAASPNKELAKEFLENYLLTDEGLEAVNKDKPLGAVALKSY
EEELAKDPRIAATMENAQKGEIMPNIPQMSAFWYAVRTAVINAASGRQTVDEALKDAQTNSGSD
ITSLYKKAEGGTENLYFQGHTERRVPFSLLRGPSWDPFRDWYPHSRLFDQAFGLPRLPEEWS
QWLGGSSWPGYVRPLPPAAIESPAVAAPAYSRALSRQLSCLSYDTEILTVEYGFLPIGKIVEERIE
CTVYTVDKNGFVYTQPIAQWHNRGEQEVFEYCLEDGSIIRATKDHKFMTTDGQMLPIDEIFERG
LDLKQVDGLPHHHHHH

*6xHis-$Cfa^C$-HSPB1[S83C-ACD-CTD]:*
MKSSHHHHHHVKIISRKSLGTQNVYDIGVGEPHNFLLKNGLVASNCGVSEIRHTADRWRVSLDV
NHFAPDELTVKTKDGVVEITGKHEERQDEHGYISRCFTRKYTLPPGVDPTQVSSSLSPEGTLTV
EAPMPKLATQSNEITIPVTFESRAQLGGPEAAKSDETAAK

*HSPB1-GSKCK:*
MTERRVPFSLLRGPSWDPFRDWYPHSRLFDQAFGLPRLPEEWSQWLGGSSWPGYVRPLPPA
AIESPAVAAPAYSRALSRQLSSGVSEIRHTADRWRVSLDVNHFAPDELTVKTKDGVVEITGKHEE
RQDEHGYISRCFTRKYTLPPGVDPTQVSSSLSPEGTLTVEAPMPKLATQSNEITIPVTFESRAQL
GGPEAAKSDETAAKGSKCK

HSPB1 was prepared in Rosetta (DE3) competent *Escherichia coli* cells (MilliporeSigma, Burlington, MA) that had been transformed with the appropriate HSPB1 plasmid. Seed cultures were grown to saturation from freshly transformed colonies and inoculated at a 1% v/v ratio into Luria-Bertani medium supplemented with kanamycin (50 µg/mL), or M9 medium supplemented with kanamycin (50 µg/mL), $^{13}C_6$-glucose (2 g/L) and $^{15}N$-enriched ammonium chloride (1 g/L). Cultures were grown in LB at 37°C to an OD600 of ~ 0.6, and protein expression was induced by the addition of 1 mM isopropyl-β-D-thiogalactoside. Cultures were incubated for 4 hours at 37˚C before being harvested by centrifugation at 5,000 xg at 4°C for 30 minutes. After decanting the supernatant, cell pellets were either immediately used for purification of HSPB1 or stored at -80°C for later use.

**Purification of full-length and GSKCK-tagged HSPB1**

Full-length and GSKCK-tagged HSPB1 were purified using the ammonium sulfate precipitation method described in Ref. (31-33) with some modifications. This method enabled the preparation of HSPB1 without the need for a solubility or affinity tag. Briefly, the HSPB1 cell pellet was thawed if necessary, resuspended in lysis buffer (50 mM sodium chloride, 50 mM sodium phosphate, Roche cOmplete$^{TM}$ EDTA-free Protease Inhibitor Cocktail, pH 7.4) and lysed by pulsed sonication for 30 minutes using a Qsonica sonicator with a 1/8” diameter probe tip at 12 kHz (60%) output and 4°C. The lysate was cleared by centrifugation at 30,000 xg for 30 minutes and the supernatant was decanted into an Erlenmeyer flask with a stir bar. Saturated ammonium sulfate (4.1 M) was added dropwise to this solution, to a final concentration of 40% saturation (1.64 M) at room temperature. The solution was allowed to stir for 30 minutes, after which the precipitated protein was pelleted by centrifugation at 30,000 xg for 30 minutes at 4 °C. The supernatant was discarded, and the pellet was redissolved in 20 mM tris(hydroxymethyl)aminomethane, 10 mM magnesium chloride, 30 mM ammonium chloride, pH 7.6 at 25 ˚C. The resulting solution was desalted by running over PD-10 desalting columns packed with Sephadex G-25 resin (Cytiva, Marlborough, MA), equilibrated in the same buffer. The desalted HSPB1 solution was further purified by ion exchange chromatography over a DEAE Sepharose FF anion exchange column (Cytiva, Marlborough, MA), equilibrated in the same buffer. Protein was eluted from the column using a 0-250 mM sodium chloride gradient. The fractions containing HSPB1 were concentrated and subjected to reverse-phase HPLC purification over a Waters XBridge Peptide BEH C18 OBD prep column (130 Å pore size, 10 µm particle size, 19 mm X 250 mm) using a Waters 2535 Binary Gradient Module equipped with a 2484 UV/Vis detector (Waters Corporation, Milford, MA). Sample purity was confirmed using an analytical Waters XBridge Peptide BEH C18 column (300 Å pore size, 5 µm particle size, 2.1 mm X 100 mm) on the same HPLC system. Sample identity was confirmed by intact mass TOFMS using an Agilent 1260 Infinity Binary LC coupled to a 6230 TOFMS system (Agilent Technologies, Santa Clara, CA). All HPLC purifications utilized a gradient method of water and acetonitrile with 0.1% trifluoroacetic acid as the mobile phases. After HPLC purification, samples were lyophilized and refolded by dialysis at 4 °C from 6 M guanidine hydrochloride, 150 mM sodium chloride, 50 mM sodium phosphate, pH 7.4, into the same buffer without denaturant. If samples were not used immediately after preparation, they were concentrated to ~1 mM and flash frozen at -80 °C for storage.

**Mass photometry**
A Refeyn TwoMP mass photometer (Refeyn Ltd., UK) was used to analyze the HSPB1 samples. Sample stocks were prepared at low µM concentrations for full-length HSPB1 and FUS LC in a 150 mM sodium chloride, 50 mM sodium phosphate, pH 7.4 buffer, except in the case where salt concentration was screened. Prior to data acquisition, Beta-amylase was used as a protein calibrant with molecular weights set to 56 (monomer), 112 (dimer), and 224 (tetramer) kDa for distinct oligomeric species detected. Data was then acquired first by using 10 µL of buffer to calibrate and focus the instrument, then followed by the addition of 10 µL of protein sample at a 2X concentration compared to the final desired concentration. Final concentrations varied by experiment but were in the range of 250 to 1000 nM for full-length HSPB1 and 500 nM to 5000 nM for FUS LC. Gaussian curve-fitting was applied using Refeyn DiscoverMP software to determine the molecular weight and distribution of oligomers in the sample.

**Analytical Ultracentrifugation**
The analytical ultracentrifugation sedimentation velocity experiment was performed using a ProteomeLab XL-I (Beckman Coulter, Brea, CA) analytical ultracentrifuge, with absorbance detection at 280 nm. Samples of full-length HSPB1 were prepared in 50 mM sodium chloride, 50 mM sodium phosphate, +/- 2 mM tris(2-carboxyethyl)phosphine (TCEP), pH 7.4 at 0.3 mg/ml (A280 = 0.5) and loaded into 2-channel cells equipped with sapphire windows and spun using an An-50 Ti 8-place rotor at 40,000 rpm and 20°C for 20 hours. Data were analyzed using Sedfit (34).

**Negative Staining and TEM**

Freshly prepared glow-discharged Formvar 300 mesh carbon-on-copper grids (Electron Microscopy Sciences, Hatfield, PA) were placed onto 10 µl drops of sample arrayed on a Parafilm sheet. After allowing protein to adsorb onto the grids for 5 minutes, the grids were washed with buffer 3 times and applied to drops of 1% w/v uranyl acetate (Ladd Research Industries, Williston, VT) arrayed on a Parafilm sheet and allowed to stain for 1 minute. After staining, excess uranyl acetate solution was wicked from each grid and the grids were allowed to dry for at least an hour before insertion into the microscope. Grids were viewed using a JEOL JEM-1400Plus transmission electron microscope operating at 80 kV and micrographs were recorded using a Gatan OneView digital camera.

**Purification and splicing of HSPB1 intein constructs**

All intein constructs included a 6x-His tag and were purified by the same Ni-NTA affinity chromatography approach. Briefly, the cell pellet was thawed if necessary, resuspended in lysis buffer (200 mM sodium chloride, 50 mM sodium phosphate, 2 mM TCEP, Roche cOmpleteTM EDTA-free Protease Inhibitor Cocktail, pH 8.0) and lysed by pulsed sonication for 30 minutes using a Qsonica sonicator with a 1/8" diameter probe tip at 12 kHz (60%) output and 4°C. The lysate was cleared by centrifugation at 30,000 RCF for 30 minutes and the supernatant was incubated with 3 mL Thermo Scientific HisPur Ni-NTA Resin (Thermo Fisher Scientific, Waltham, MA) per liter of culture for 30 min at 4°C. The suspension of beads was washed with 10 column volumes of lysis buffer containing 10 mM imidazole, and protein was eluted with two column volumes of lysis buffer containing 250 mM imidazole. Ni-NTA binding was confirmed using SDS-PAGE gel electrophoresis. Immediately after elution, the fractions containing the eluted protein were combined and dialyzed into splicing buffer (200 mM sodium chloride, 50 mM sodium phosphate, 2 mM TCEP, 1 mM EDTA, pH 8.0) at 25°C for 4 hours, with a buffer exchange after 2 hours.

To prepare segmentally labeled constructs in which the NTD is $^{15}$N, $^{13}$C-labeled ([$^{15}$N,$^{13}$C-NTD]-ACD-CTD), the $^{15}$N,$^{13}$C-NTD-6xHis-MBP-TEV-HSPB1NTD-Cfa$^{N}$-6xHis and 6xHis-Cfa$^{C}$-HSPB1ACD-CTD samples were diluted in splicing buffer post-dialysis and incubated for 15 minutes at 25°C before mixing to a final concentration of 20 µM $^{15}$N,$^{13}$C-6xHis-MBP-TEV-HSPB1NTD-Cfa$^{N}$-6xHis and 30 µM 6xHis-Cfa$^{C}$-HSPB1ACD-CTD at a total reaction volume of 180 mL. The intein splicing reaction was allowed to proceed overnight at 25°C. To prepare segmentally labeled HSPB1 in which the ACD and CTD are $^{15}$N, $^{13}$C-labeled, (NTD-[$^{15}$N,$^{13}$C-ACD-CTD]), the same protocol was followed where 6xHis-MBP-TEV-HSPB1NTD-Cfa$^{N}$-6xHis and $^{15}$N,$^{13}$C-6xHis-Cfa$^{C}$-HSPB1ACD-CTD were mixed to final protein concentrations of 20 µM and 30 µM respectively, and allowed to react overnight at 25˚C. After splicing, the reaction was incubated with 6xHis-tagged TEV protease at a molar ratio of 1:50 TEV to protein for 4 hours at 25°C to remove the 6xHis-MBP tag from the spliced product. Splicing and cleavage were monitored by SDS-PAGE and analytical HPLC (**Fig. S6**). 6 M guanidine hydrocholoride, 5 mM TCEP, and 0.1% TFA was added to the TEV splicing mixture and the sample was purified using reverse-phase HPLC over a Waters XBridge Peptide BEH C18 OBD prep column (130 Å pore size, 10 µm particle size, 19 mm X 250 mm) using a Waters 2545 Binary Gradient Module equipped with a 2484 UV/Vis detector (Waters Corporation, Milford, MA). Sample purity was confirmed using an analytical Waters XBridge Peptide BEH C18 column (300 Å pore size, 5 µm particle size, 2.1 mm X 100 mm) on the same HPLC system. Sample identity was confirmed by intact mass QTOFMS using an Agilent 1260 Infinity Binary LC coupled to a 6230 TOFMS system (Agilent Technologies, Santa Clara, CA). All HPLC purifications utilized a gradient method of water and acetonitrile with 0.1% trifluoroacetic acid as the mobile phase. After HPLC purification, samples were lyophilized and refolded by dialysis from 6 M guanidine hydrochloride, 150 mM sodium chloride, 50 mM sodium phosphate, and 2mM TCEP, pH 7.4, into the same buffer but without denaturant. If samples were not used immediately after preparation, they were concentrated, and flash frozen at -80 °C for storage.

**Expression of FUS LC**

6xHis-MBP-TEV-FUS(1-160) and 6xHis-MBP-TEV-FUS(1-160)-CSG (used for fluorophore attachment) were expressed and purified as described previously (35, 36). In summary, FUS LC seed cultures were grown overnight and inoculated at a 1% v/v ratio into Luria-Bertani medium supplemented with kanamycin (50 mg/mL). The cultures were then grown at 37 ˚C to an OD600 of 0.7 when protein expression was induced by the addition of 1 mM isopropyl-b-D-thiogalactoside. Cultures were then grown for an additional 4 hours at 37 ˚C before being harvested for purification.

*6xHis-MBP-TEV-FUS(1-160):*
MGSDKIHHHHHHSSGTKIEEGKLVIWINGDKGYNGLAEVGKKFEKDTGIKVTVEHPDKLEEKFP
QVAATGDGPDIIFWAHDRFGGYAQSGLLAEITPDKAFQDKLYPFTWDAVRYNGKLIAYPIAVEAL
SLIYNKDLLPNPPKTWEEIPALDKELKAKGKSALMFNLQEPYFTWPLIAADGGYAFKYENGKYDI
KDVGVDNAGAKAGLTFLVDLIKNKHMNADTDYSIAEAAFNKGETAMTINGPWAWSNIDTSKVNY
GVTVLPTFKGQPSKPFVGVLSAGINAASPNKELAKEFLENYLLTDEGLEAVNKDKPLGAVALKSY
EEELAKDPRIAATMENAQKGEIMPNIPQMSAFWYAVRTAVINAASGRQTVDEALKDAQTNSGSD
ITSLYKKAEGGTENLYFQGMASNDYTQQATQSYGAYPTQPGQGYSQQSSQPYGQQSYSGYS
QSTDTSGYGQSSYSSYGQSQNTGYGTQSTPQGYGSTGGYGSSQSSQSSYGQQSSYPGYGQ
QPAPSSTSGSYGSSSQSSSYGQPQSGSYSQQPSYGGQQQSYGQQQSYNPPQGYGQQNQY
NS

*6xHis-MBP-TEV-FUS(1-160)-CSG:*
MGSDKIHHHHHHSSGTKIEEGKLVIWINGDKGYNGLAEVGKKFEKDTGIKVTVEHPDKLEEKFP
QVAATGDGPDIIFWAHDRFGGYAQSGLLAEITPDKAFQDKLYPFTWDAVRYNGKLIAYPIAVEAL
SLIYNKDLLPNPPKTWEEIPALDKELKAKGKSALMFNLQEPYFTWPLIAADGGYAFKYENGKYDI
KDVGVDNAGAKAGLTFLVDLIKNKHMNADTDYSIAEAAFNKGETAMTINGPWAWSNIDTSKVNY
GVTVLPTFKGQPSKPFVGVLSAGINAASPNKELAKEFLENYLLTDEGLEAVNKDKPLGAVALKSY
EEELAKDPRIAATMENAQKGEIMPNIPQMSAFWYAVRTAVINAASGRQTVDEALKDAQTNSGSD
ITSLYKKAEGGTENLYFQGMASNDYTQQATQSYGAYPTQPGQGYSQQSSQPYGQQSYSGYS
QSTDTSGYGQSSYSSYGQSQNTGYGTQSTPQGYGSTGGYGSSQSSQSSYGQQSSYPGYGQ
QPAPSSTSGSYGSSSQSSSYGQPQSGSYSQQPSYGGQQQSYGQQQSYNPPQGYGQQNQY
NSCSG

**Purification of FUS LC**

For purification, the cell pellet was resuspended in lysis buffer (300 mM sodium chloride, 20 mM sodium phosphate, Roche cOmplete EDTA-free Protease Inhibitor Cocktail (MilliporeSigma, Burlington, MA), pH 7.4), and lysed by pulsed sonication for 30 minutes using a Qsonica sonicator with a 1/8” diameter probe tip at 12 kHz (60%) output and 4°C before centrifugation to clear the lysate. The supernatant was then incubated with Thermo Scientific HisPur Ni-NTA Resin (Thermo Fisher Scientific, Waltham, MA) for 30 minutes at 4 ˚C and the protein was eluted with a 250 mM imidazole. The eluted protein was incubated with TEV protease at a molar ratio of 1:50 TEV to protein for 5 hours at 25 ˚C for tag cleavage and then 8 M urea was added to the mixture to solubilize the aggregated FUS LC. The mixture was then diluted into 20 mM CAPS (pH 11) and 150 mM sodium chloride and subjected to size-exclusion chromatography over a GE HiLoad 16/600 Superdex 75-pg column (GE Healthcare, Chicago, IL). Purity was verified by HPLC and mass spectrometry. FUS LC-CSG was subjected to the same purification protocol except labeling with Cy3-malemide (see below) was introduced after the Ni-NTA purification, followed byTEV cleavage and size-exclusion chromatography.

**Insulin aggregation assay**

To characterize the chaperone activity of HSPB1, an insulin aggregation assay was performed (37). A 250 µM stock solution of insulin from bovine pancreas (MilliporeSigma, Burlington, MA) was prepared by first dissolving 1.5 mg of insulin in 100 µL of 0.05 M HCl, then diluting the solution in 900 µL of buffer (150 mM sodium chloride, 50 mM sodium phosphate, pH 7.4). An extinction coefficient of 6080 $M^{-1}cm^{-1}$ at A280 nm was used to determine the insulin concentration. Protein samples for the assay were prepared in 96-well plates, where the total assay volume was 100 µL. HSPB1 was added at concentrations ranging from 0 to 10 µM (**Fig. S4**) while the insulin concentration was kept constant at 50 µM in 150 mM sodium chloride, 50 mM sodium phosphate, pH 7.4 buffer. The assay plate was incubated at 37 ˚C for 5 minutes before the addition of 20 mM DTT to induce insulin aggregation. Following DTT addition, light scattering at A360 nm was measured in 30 s intervals for 45 min with shaking before measurement. Data was normalized to peak insulin aggregation (based upon highest A360 intensity) in the absence of HSPB1.

**Fluorescent labeling and microscopy of proteins**

Cy5-labeled HSPB1 was produced by using an HSPB1-GSKCK construct which was expressed and purified as described above with some modifications. We used this construct to avoid labeling the only native cysteine residue Cys 137 in the ACD domain, which can be involved in structurally important intermolecular disulfide bond. After ammonium sulfate precipitation, the protein pellet was resuspended in reaction buffer (150 mM sodium chloride, 50 mM sodium phosphate, 1 mM EDTA, 2mM TCEP, pH 7.4). Excess Cy5-malemide (Lumiprobe, Hunt Valley, MD) was added to the mixture and the reaction was allowed to proceed for 2 hours at 25 ˚C in the dark before quenching with excess β-mercaptoethanol (>200 M eq.). The resulting mixture was desalted by running over PD-10 desalting columns packed with Sephadex G-25 resin (Cytiva, Marlborough, MA) equilibrated in the same buffer. The sample was further purified by reverse-phase HPLC over a Waters XBridge Peptide BEH C18 OBD prep column (300 Å pore size, 5 µm particle size, 10 mm X 250 mm) using a Waters 2545 Binary Gradient Module equipped with a 2484 UV/Vis detector (Waters Corporation, Milford, MA). All HPLC purifications utilized a gradient method of water and acetonitrile with 0.1% trifluoroacetic acid as the mobile phases. Sample identity and single cysteine-labeled product was confirmed by intact mass TOFMS using an Agilent 1260 Infinity Binary LC coupled to a 6230 TOFMS system (Agilent Technologies, Santa Clara, CA). The final purified protein was 100% labeled.

Cy3-labeled FUS LC was produced as described previously (35). Briefly, Ni-NTA-purified FUS LC-CSG was diluted to a concentration of 20 µM in reaction buffer (300 mM sodium chloride, 20 mM sodium phosphate, pH 7.4) and 500 µM TCEP and 80 µM Cy3-maleamide (4 M eq.; APExBIO, Houston, TX) were added. The reaction was allowed to proceed for 60 s at 25°C in the dark before quenching with excess β-mercaptoethanol (>200 M eq.) and dialysis into 300 mM sodium chloride, 20 mM sodium phosphate, and 100 µM TCEP (pH 7.4) at 4°C. The labeled protein was then subjected to TEV cleavage and size exclusion chromatography as described above. Labeling efficiency was determined to be 33% by analytical HPLC and mass spectrometry.

FUS LC samples for microscopy were prepared starting with FUS LC (including 5% FUS LC-CSG-Cy3) stock in 150 mM sodium chloride, 20 mM CAPS, pH 11. The stock was then diluted in a buffer containing 150 mM sodium chloride, 50 mM sodium phosphate (pH 7.4) at 4˚C to induce phase separation. HSPB1 (including 5% HSPB1-GSKCK-Cy5) was added concurrently to prepare a 5:1 molar ratio (300:60 µM) of FUS LC:HSPB1 sample. The phase-separated solution was then added to a µ-Slide 18-well glass bottom multi-well plate (Ibidi) for imaging with a Leica SP8 microscope with a 60X oil immersion objective.

**Magic angle spinning NMR experiments**

MAS experiments at moderate spinning frequencies were performed using 3.2-mm thin-walled zirconia MAS rotors with 50 µL sample volume. Spectra were acquired on a 750-MHz (17.6 T) NMR spectrometer equipped with a 3.2-mm $E^{free}$ triple resonance HCN MAS probe (Bruker Biospin, Billerica, MA). All experiments were performed at an MAS frequency of 11.111 kHz. Samples were cooled with a stream of dry nitrogen gas maintained at 275 or 295 K while we estimate that the sample temperature during the MAS experiments is 10–15° higher. MAS NMR experimental parameters can be found in the supporting materials and methods (**Tables S1-2**). Spectra were referenced using the 40.48 ppm signal of adamantane (38). Standard data processing was performed with TopSpin 4.4.0 and the data were visualized using matplotlib and nmrglue python packages (39, 40). Statistical analysis of NMR chemical shifts was performed as described previously (35).

$T_{1\rho}$ experiments were performed using 3.2-mm thin-walled zirconia MAS rotors with 50 µL sample volume. Measurements were done on segmentally labeled samples of HSPB1 alone or in the presence of phase-separated FUS LC. Briefly, segmentally labeled HSPB1 was mixed with FUS LC in 150 mM sodium chloride, 50 mM sodium phosphate, 2 mM TCEP, pH 7.2-7.4 at 4˚C to induce FUS LC phase separation. The mixture was incubated at 4˚C for 30 min before light centrifugation into a 3.2 mm thin-walled zirconia rotor to pack the phase-separated material. We estimate that the rotors packed with phase-separated mixtures of FUS LC and HSPB1 contain a ratio of 5:1 FUS LC:HSPB1. Spectra were acquired on a 750 MHz (17.6 T) NMR spectrometer equipped with a 3.2-mm $E^{free}$ triple resonance HCN MAS probe (Bruker Biospin, Billerica, MA). All experiments were performed at an MAS frequency of 11.111 kHz. Standard data processing was performed with TopSpin 4.4.0 and the data was fit to a biexponential decay function (Eq. 1) using a custom-built script in python, where $t_{SL}$ is the spin lock time (ms), and *A1* and *A2* are weighted factors to determine the signal intensity (*I*) contribution of each relaxation component.

$$\boldsymbol{I(t_{SL}) = A1e^{\left(-\frac{t_{SL}}{T_{1\rho,1}}\right)} + A2e^{\left(-\frac{t_{SL}}{T_{2\rho,2}}\right)}} \quad \textbf{(Eq. 1)}$$

Fast $^{1}$H-based MAS experiments were performed using a 0.4-mm zirconia MAS rotor with a 0.12 µL sample volume (~ 25 µg of protein). Spectra were acquired on an 800 MHz (18.8 T) Bruker Avance Neo Spectrometer equipped with Bruker standard bore 0.4-mm triple resonance HCN CPMAS probe (Bruker Biospin, Ettlingen, Germany). All experiments were performed at an MAS frequency of 160 kHz. Samples were cooled by controlling the temperature of the bearing gas, we estimate that the sample temperature during the MAS experiments is 25˚C. MAS NMR experimental parameters can be found in the supporting materials and methods (**Table S3**). Data processing was performed with TopSpin 4.4.0 and the data were apodized using the QSINE window function for both dimensions, with a sine bell shift of 3. Spectra were referenced based upon the $^{1}$H-reference spectrum of silicon-grease according to IUPAC referencing standards and subsequently converted to the DSS scale. Data were visualized using matplotlib and nmrglue python packages (39, 40). Previously reported solution NMR assignments for monomer and dimer constructs of HSPB1 were accessed through the Biological Magnetic Resonance Bank (BMRB) and overlaid using matplotlib (40, 41).

**Cryogenic electron microscopy (cryoEM) sample preparation and data collection**

Samples of HPSB1 were prepared in 50 mM sodium chloride, 50 mM sodium phosphate, 5 mM TCEP, pH 7.4 at concentrations ranging from 0.5 mg/ml to 4 mg/ml. Samples were allowed to equilibrate at room temperature for at least 1 hour before application to the grid. Freshly glow-discharged Quantifoil R 2/1 holey carbon-on-copper grids (Quantifoil Micro Tools GmbH, Jena, Germany) were loaded into a Vitrobot Mark II System (Thermo Fisher Scientific, Waltham, MA) and 4 µl of HSPB1 sample were applied at 4°C and 100% relative humidity to prevent sample evaporation. The blotting parameters were set to blot for 4 seconds with a blot force of -10. After blotting, the grid was plunge-frozen into liquid ethane. Following plunge freezing, the grids were clipped and stored under liquid nitrogen until data collection.

For data collection, grids were loaded into an autoloader cassette and transferred into a Titan Krios G3 transmission electron microscope equipped with a K2 Summit direct electron detector with a Bioquantum energy filter. Movies were acquired at a 300 kV accelerating voltage with a calibrated pixel size of 1.1 Å/pixel. The total electron dose was 64.5 *e*-/Å², fractionated over multiple frames to allow for dose-weighted motion correction during data processing.

**CryoEM microscopy image processing**

All cryoEM data were processed using CryoSPARC v4.2 (42). Image preprocessing, including patch motion correction, contrast transfer function estimation, blob picking, and initial 2D classification were performed during data acquisition with CryoSPARC Live. An initial dataset of ~3.48 M particles was picked from 6,204 curated movies using an elliptical blob picker with a 100 Å minimum radius and 300 Å maximum radius. Particles were extracted in 360 pixel boxes and Fourier cropped to 90 pixels. The resulting particle stack was pared down to ~1.34 M particles based on normalized cross correlation (NCC) score and power score cutoffs, and further curated to ~377 k particles via 2D classification. The final 2D classes were used to generate templates from which a subsequent round of template picking yielded ~4.09 M particles. This particle stack was pared to ~499 k particles using NCC score and power score cutoffs and further curated to ~424 k particles via 2D classification. These particles were used to train a Topaz model (43) using a learning rate of $3 \times 10^{-4}$ over 20 epochs. This model picked ~276 k particles. The resulting particle stack was curated to ~239 k particles via 2D classification. Reference-free 2D class averages from this final curated particle stack were used to illustrate oligomer architecture and symmetry in **Fig. 1c**.

#### Supporting Information Figures

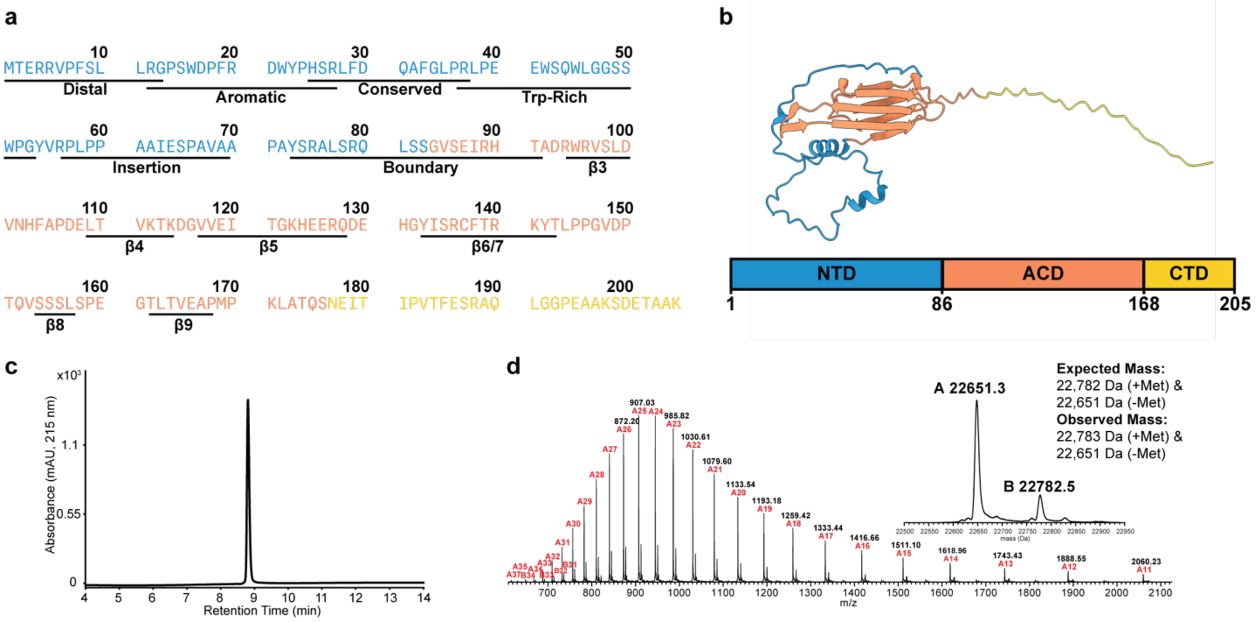


**Figure S1. (a)** Amino acid sequence of wild-type HSPB1. The NTD is shown in blue, the ACD in orange, and the CTD is in yellow. The NTD regions are annotated according to the nomenclature in Ref. (44) **(b)** Schematic of the domain organization of HSBP1 depicting a model generated with AlphaFold2 (5, 6). **(c)** Representative RP-HPLC chromatogram of HSPB1. **(c)** ESI-MS of HSPB1. The large deconvoluted peak is consistent with the molecular weight of HSPB1 lacking Met1, while the small deconvoluted peak is consistent with the molecular weight of HSPB1 that includes Met1.

a b

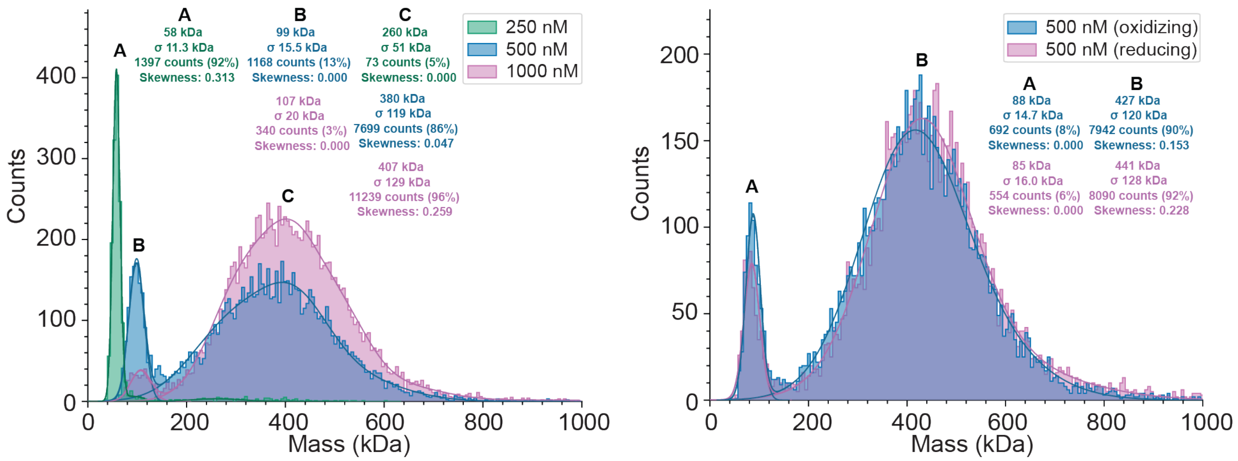


c

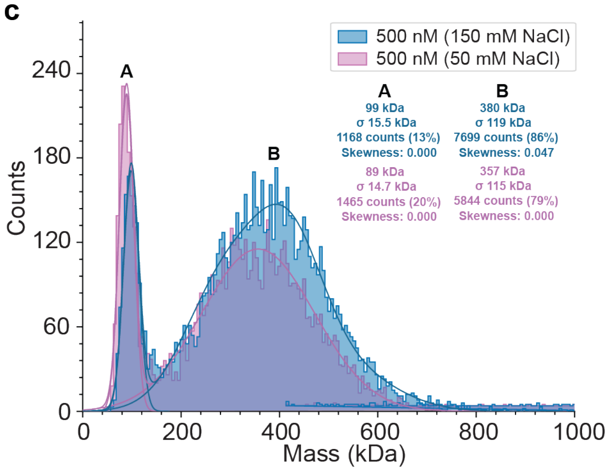


d

| Oligomer | Molecular Weight (kDa) |
|---|---|
| Monomer | 22.7(-Met) |
| Dimer | 45.3 |
| Tetramer | 90.6 |
| Hexamer | 135.9 |
| Octamer | 181.2 |
| Dodecamer | 271.8 |
| 16mer | 362.4 |
| 24mer | 543.6 |

**Figure S2. (a)** Mass photometry gaussian-fit analysis of HSPB1 at varying protein concentrations, **(b)** oxidizing versus reducing (2mM TCEP) conditions, and **(c)** two different salt concentrations. **(d)** Expected molecular weights for various HSPB1 oligomers.

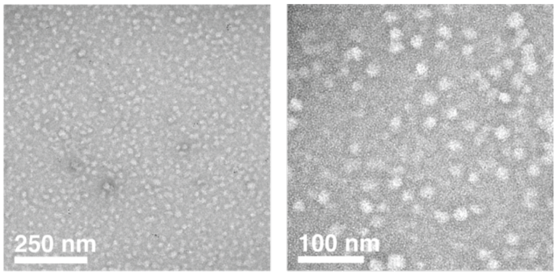


**Figure S3.** Representative negative stain TEM images of HSPB1 oligomers.

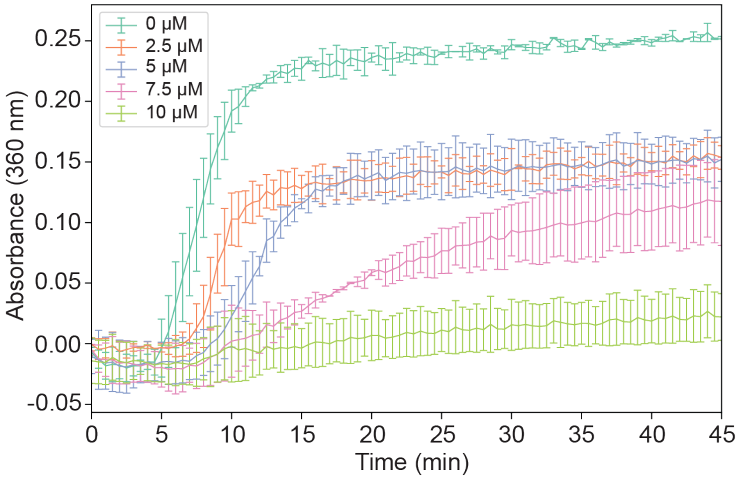


**Figure S4.** Insulin aggregation assay to test the chaperone activity of HSPB1. Insulin concentration was kept constant at 50 µM and its aggregation was induced by the addition of 20 mM DTT. A360 was measured as a function of time and in the presence of varying concentrations of HSPB1. Experiments were performed in a 150 mM NaCl and 50 mM sodium phosphate buffer (pH=7.2-7.4) at 37 ºC. Two biological replicates were used and error bars report on the standard deviation between the two replicates.

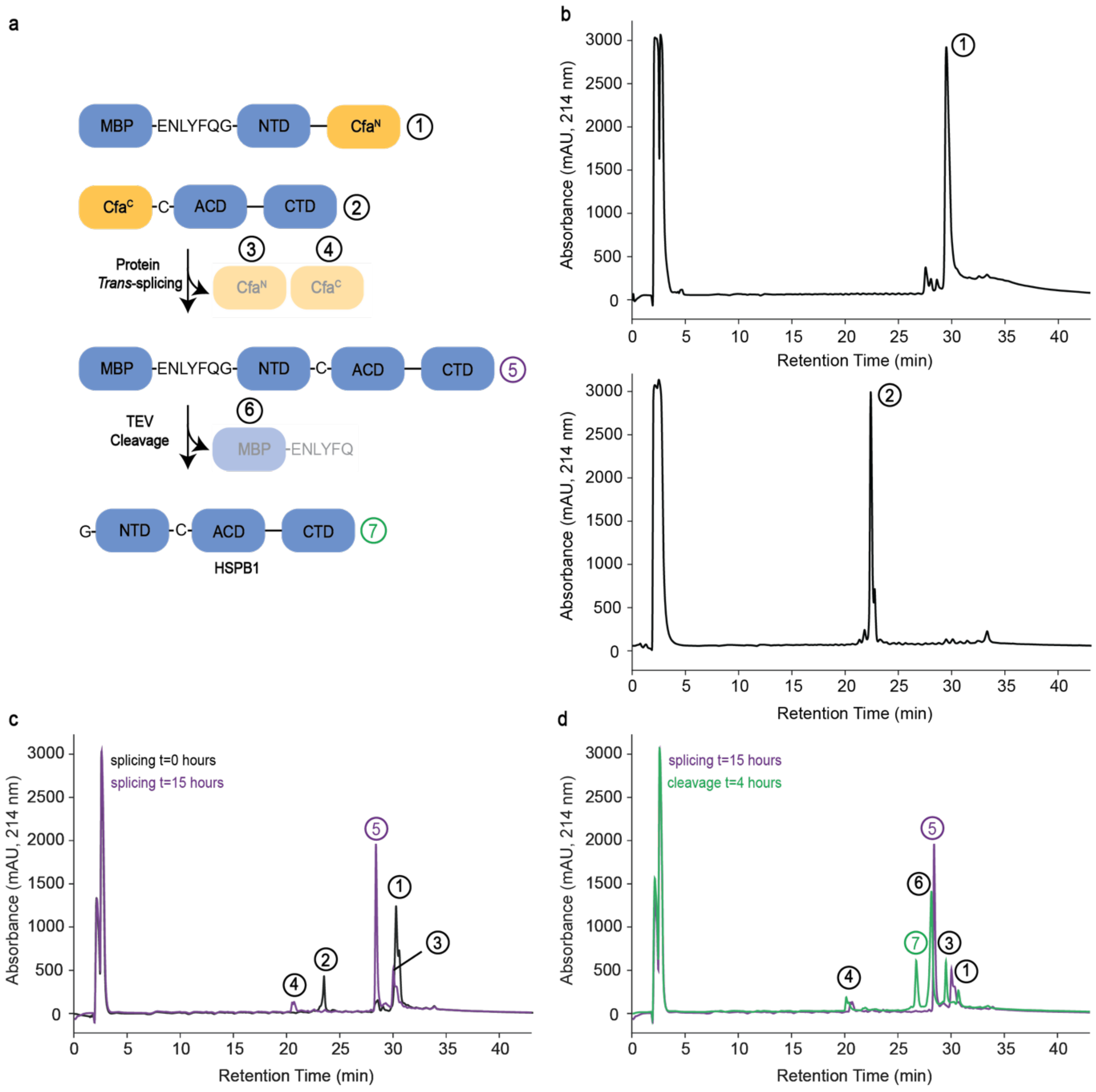


**Figure S5.** Characterization of HSPB1 splicing by RP-HPLC. **(a)** Splicing reaction and cleavage scheme. **(b)** RP-HPLC chromatograms of the starting constructs, i.e. MBP-NTD-Cfa$^N$ and Cfa$^C$-ACD-CTD. **(c)** Splicing reaction analysis by RP-HPLC. **(d)** MBP cleavage reaction analysis by RP-HPLC. All samples were analyzed on a $C_{18}$ column at a flow rate of 1mL/min. The method used started with flowing 95%/5% solvent A/B for 3 min, followed by a gradient of 30-70% solvent B from 3-43 minutes (solvent A = 0.1% TFA in $H_2O$, solvent B = 0.1% TFA in acetonitrile). See materials and methods for splicing and cleavage reaction conditions.

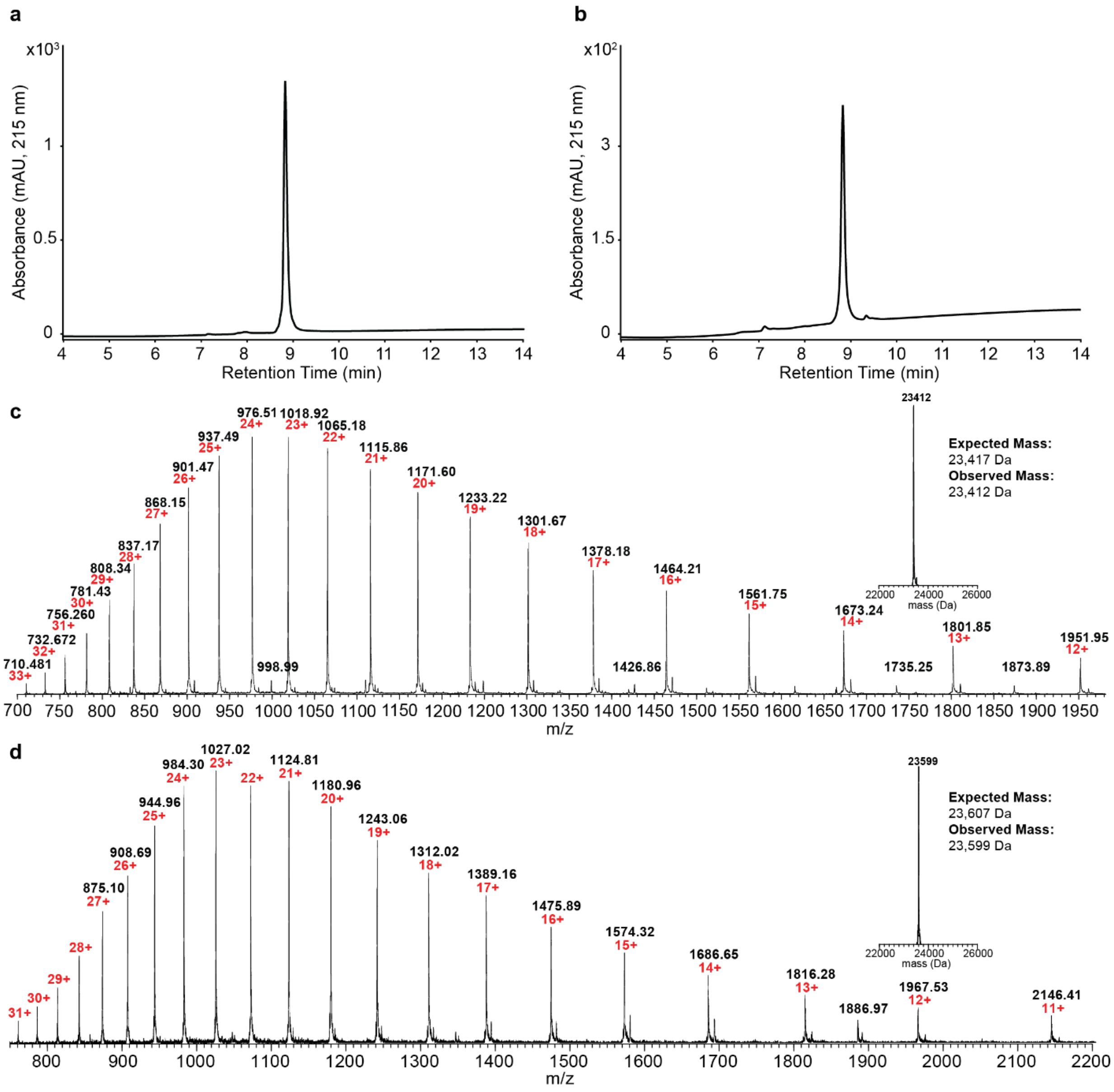


**Figure S6.** Representative RP-HPLC chromatograms and ESI-MS of the final segmentally labeled HSPB1 constructs, **(a)** and **(c)** [$^{13}C$,$^{15}N$ NTD]-ACD-CTD, and **(b)** and **(d)** NTD-[$^{13}C$,$^{15}N$ ACD-CTD].

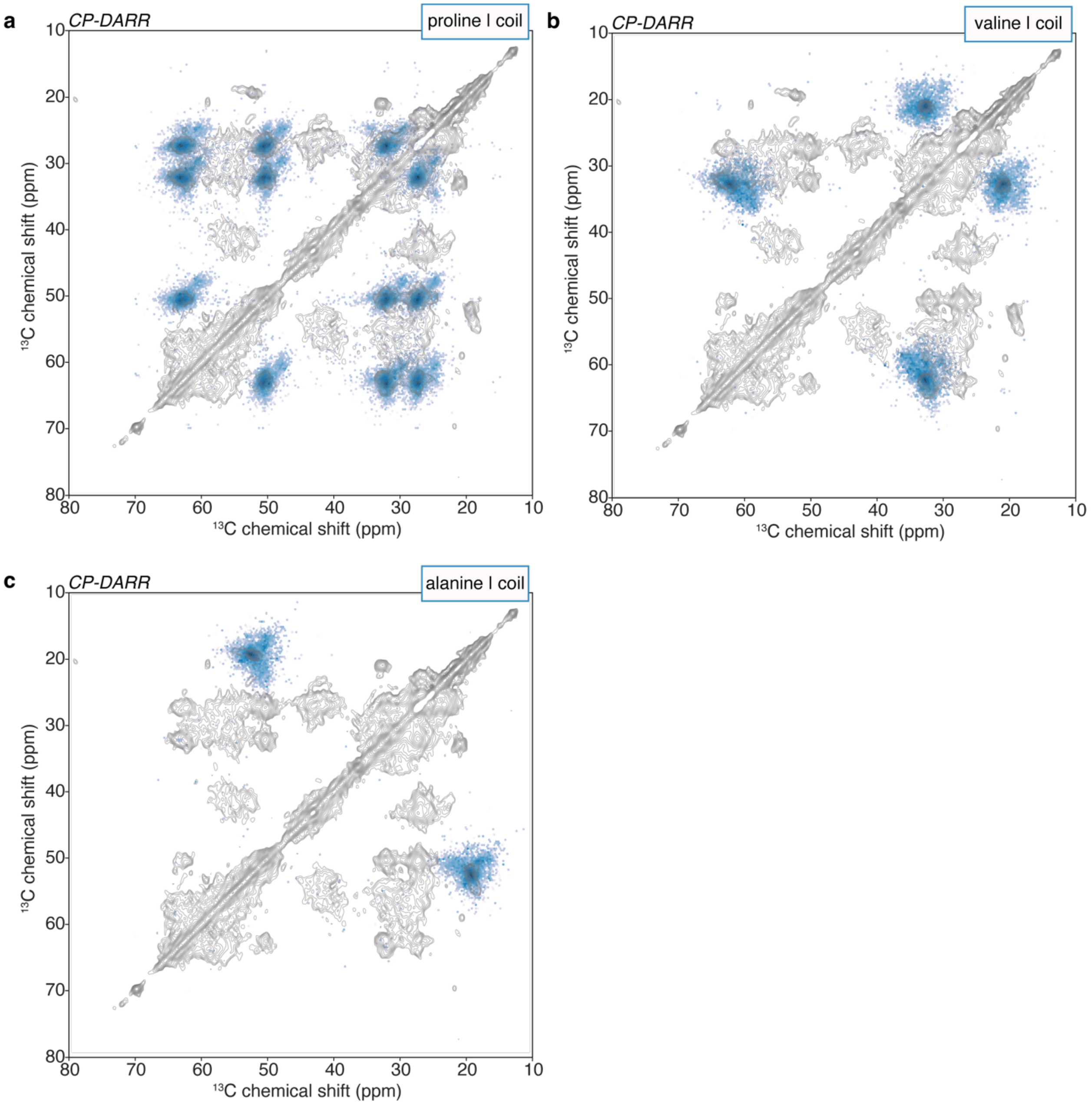


**Figure S7.** Analysis of the random coil content of the $^{13}C$-$^{13}C$ DARR correlation spectrum of NTD-labeled HSPB1 for **(a)** proline, **(b)** valine, and **(c)** alanine residues. The blue densities denote the statistical distributions of the random coil chemical shifts statistics from the BMRB.

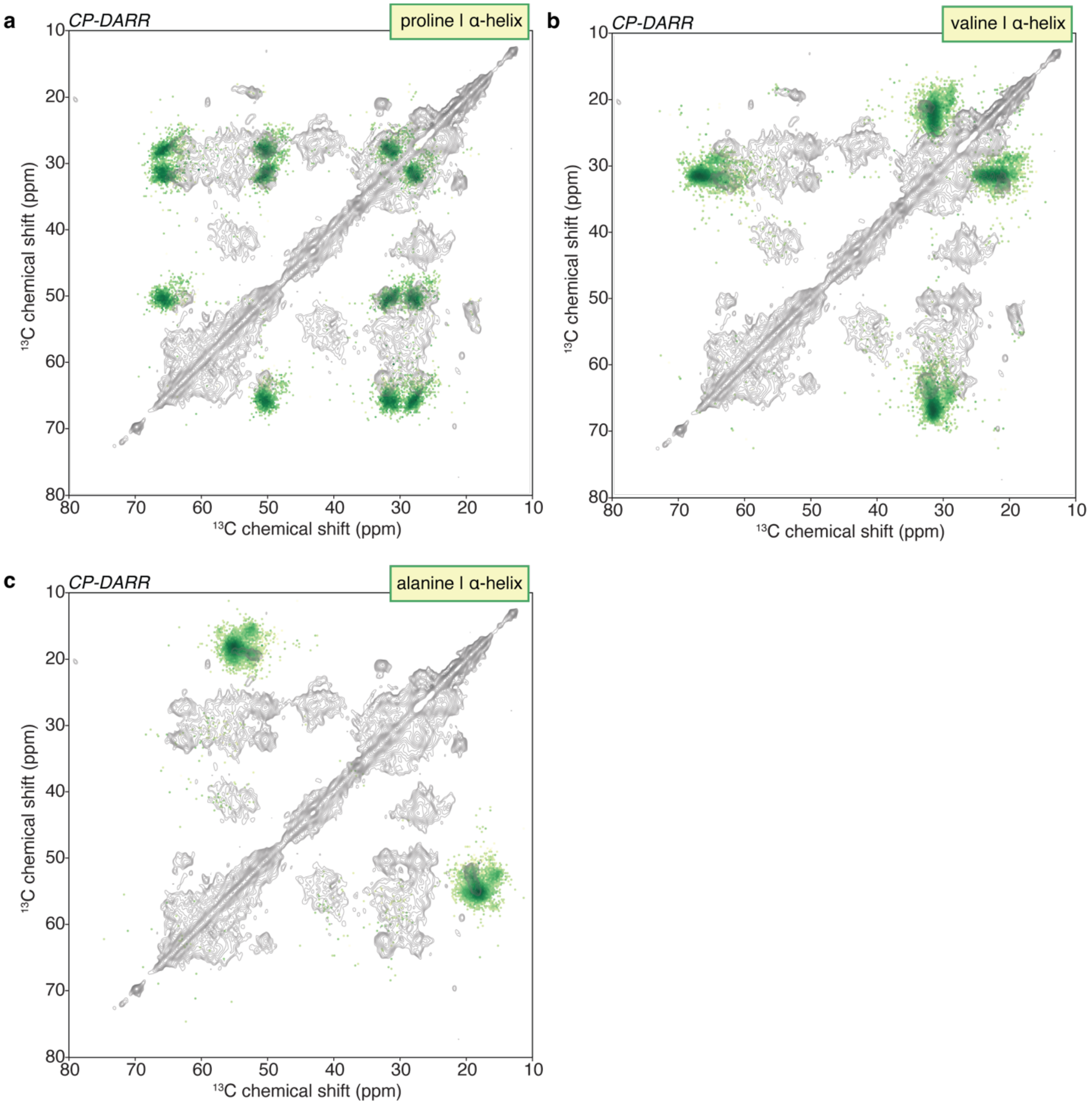


**Figure S8.** Analysis of the α-helical content of the $^{13}C$-$^{13}C$ DARR correlation spectrum of NTD-labeled HSPB1 for **(a)** proline, **(b)** valine, and **(c)** alanine residues. The green densities denote the statistical distributions of the α-helical chemical shifts statistics from the BMRB.

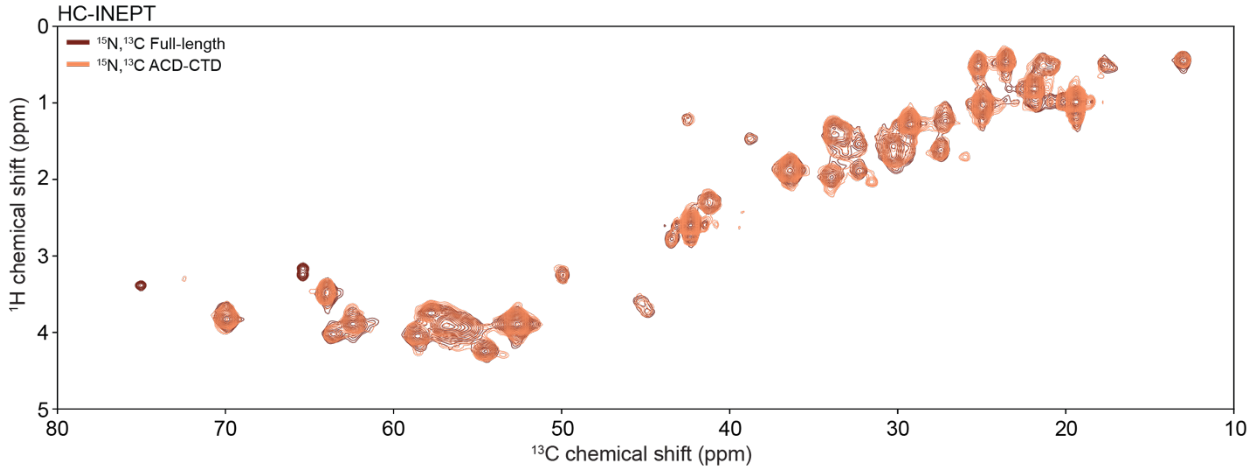


**Figure S9.** Comparison of the $^1$H-$^{13}$C INEPT spectra of fully labeled $^{13}$C,$^{15}$N HSPB1 (maroon) and segmentally labeled ACD-CTD HSPB1 constructs (orange). The two maroon cross-peaks that don't overlap with orange peaks are consistent with the $C_\alpha$ and $C_\beta$ correlations expected for Thr2 in the NTD domain.

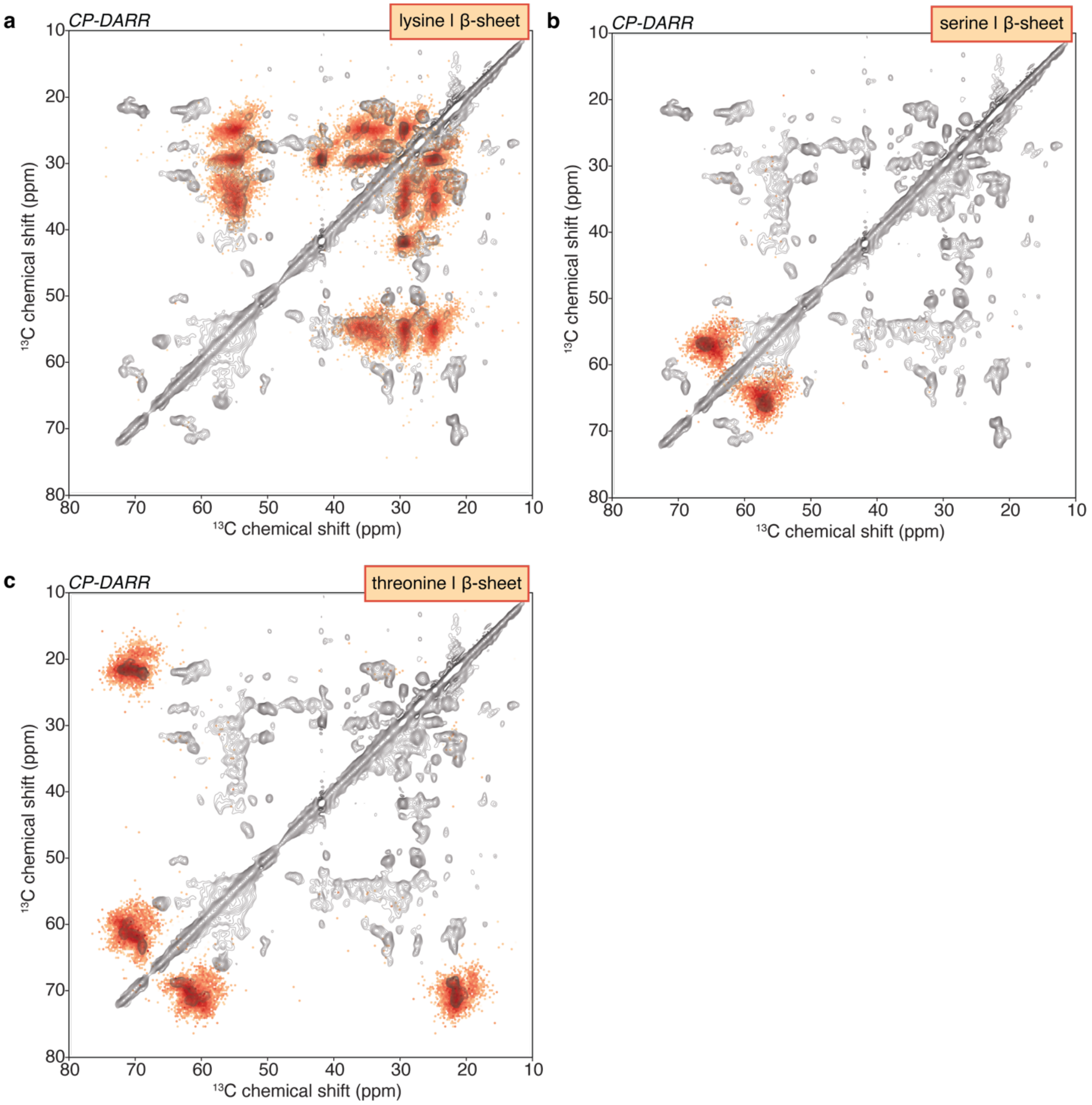


**Figure S10.** Analysis of the β-sheet content of the $^{13}C$-$^{13}C$ DARR correlation spectrum of ACD-CTD labeled HSPB1 for **(a)** lysine, **(b)** serine, and **(c)** threonine residues. The red densities denote the statistical distributions of the β-sheet chemical shifts statistics from the BMRB.

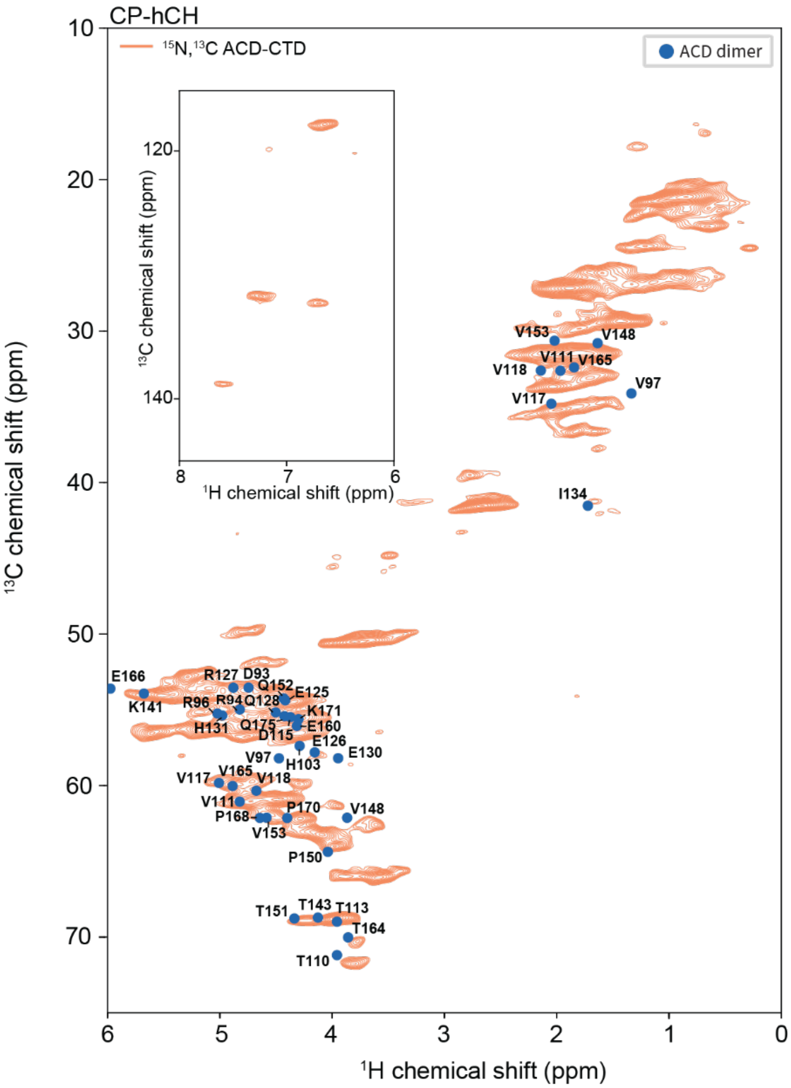


**Figure S11.** Dipolar based hCH spectrum of ACD-CTD labeled HSPB1 oligomers. The blue dots denote Hα-Cα and Hβ-Cβ solution NMR chemical shift assignments of ACD dimers (Ref.(45)). The spectrum was recorded on an 800 MHz NMR spectrometer with MAS frequency of 160 kHz.

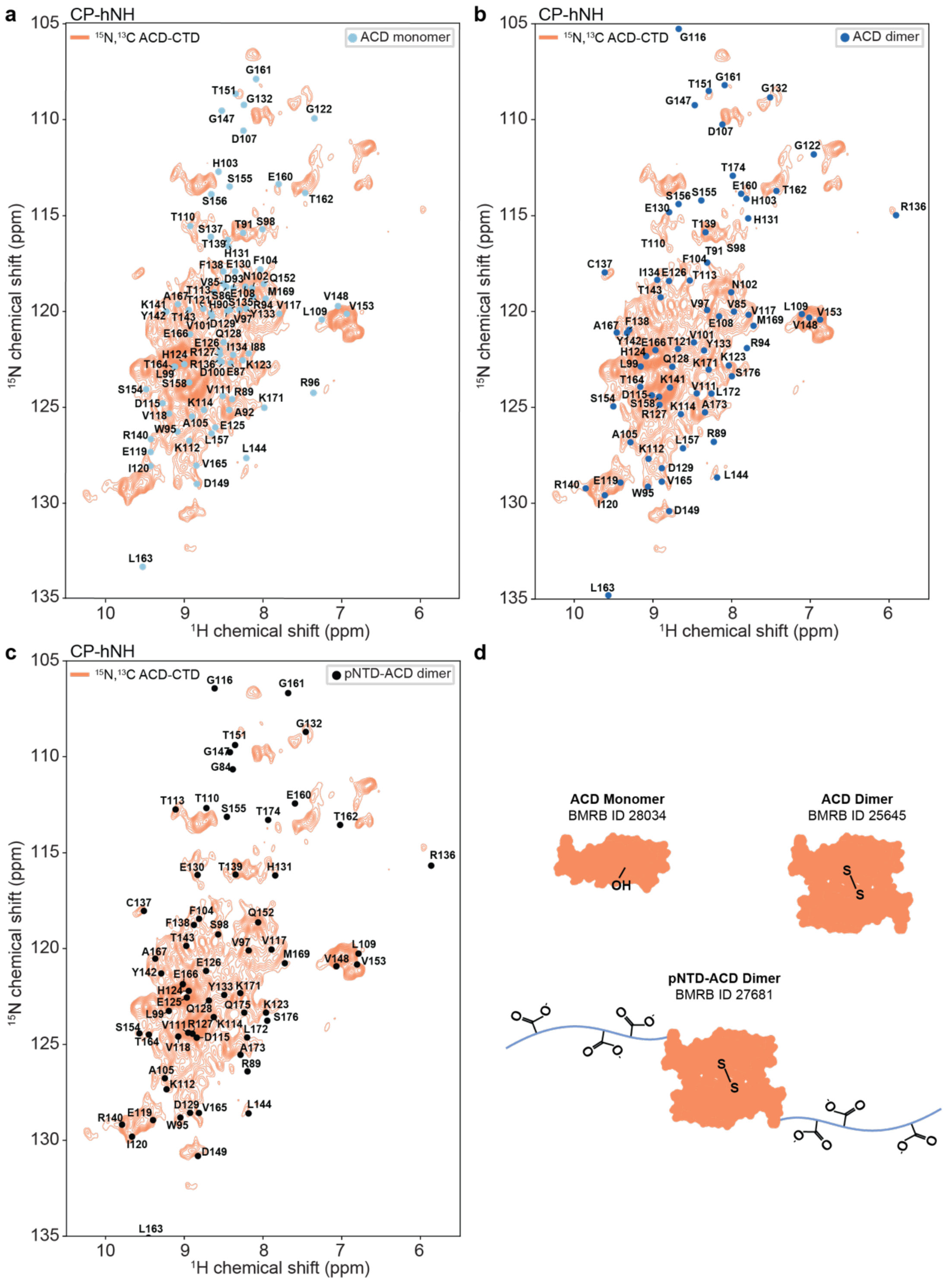


**Figure S12.** Overlays of $^{1}H$-$^{15}N$ solution NMR assignments onto the dipolar based hNH spectrum of ACD-CTD labeled HSPB1 oligomers. The assignments represent **(a)** ACD monomers (Ref. (46)), **(b)** ACD dimers (Ref.(45)) and **(c)** NTD-ACD dimers where the NTD has S15D, S78D and S82D mutations to mimic phosphorylation and promote dimerization (Ref.(44)). (d) Cartoon representation of the solution NMR constructs.

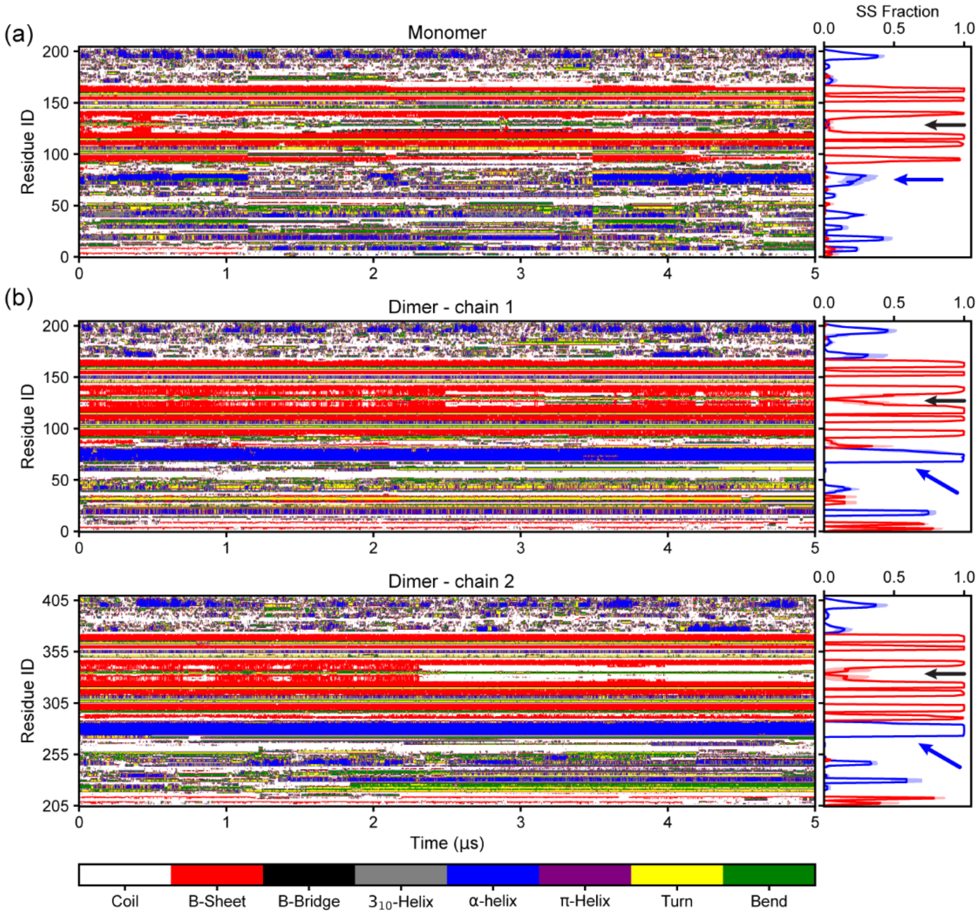


**Figure S13**. Secondary structure variation as a function of time for **(a)** an HSPB1 monomer and **(b)** each monomer in the HSPB1 dimer. Blue and black arrows mark the predicted helix region in the NTD and the loop between β6+7, respectively.

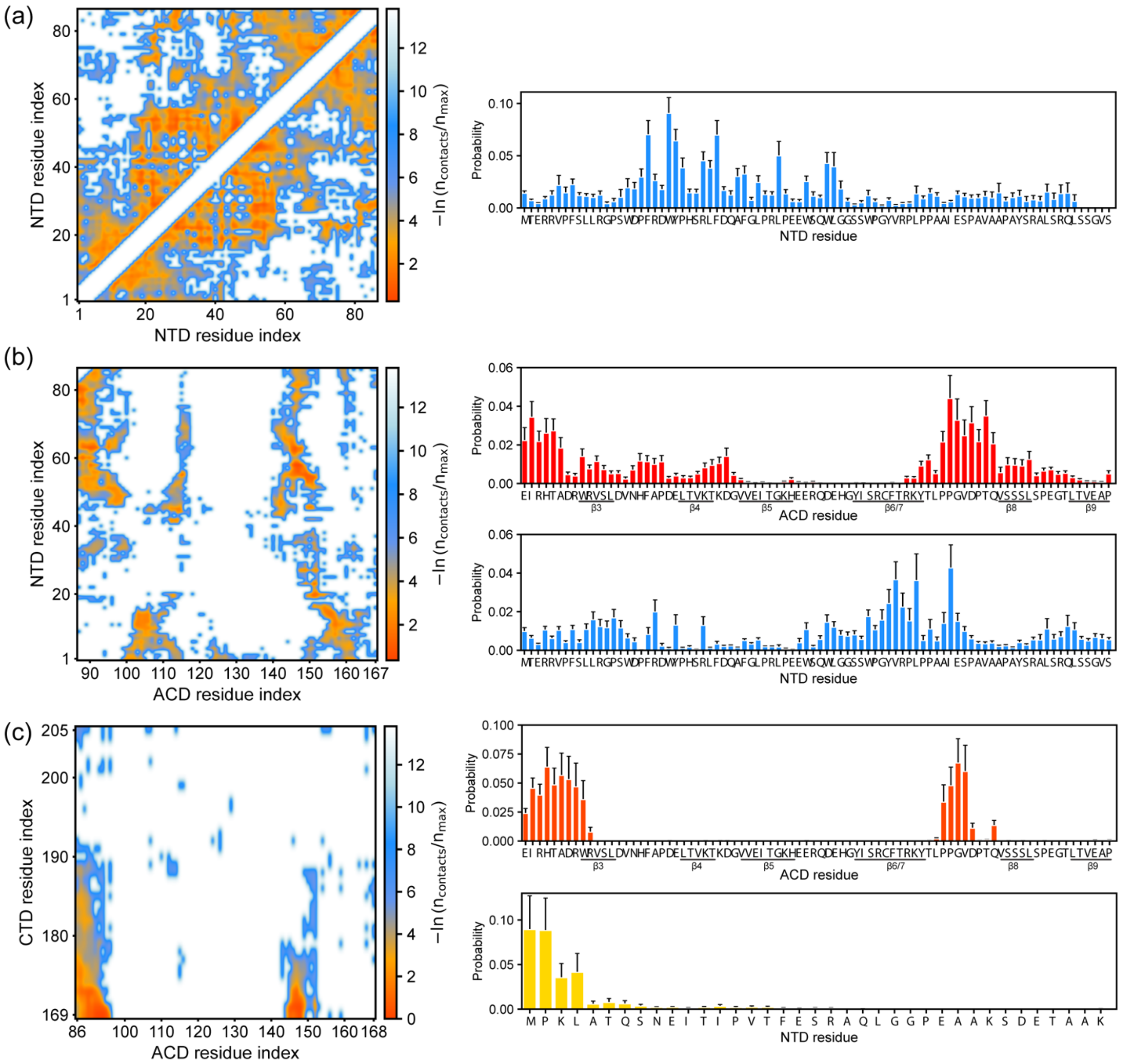


**Figure S14**. Zoomed in contact maps and residue specific contact probabilities for HSPB1 monomers from all-atom molecular dynamics simulations. **(a)** NTD-NTD contacts and, **(b)** NTD-ACD contacts, and **(c)** ACD-CTD contacts.

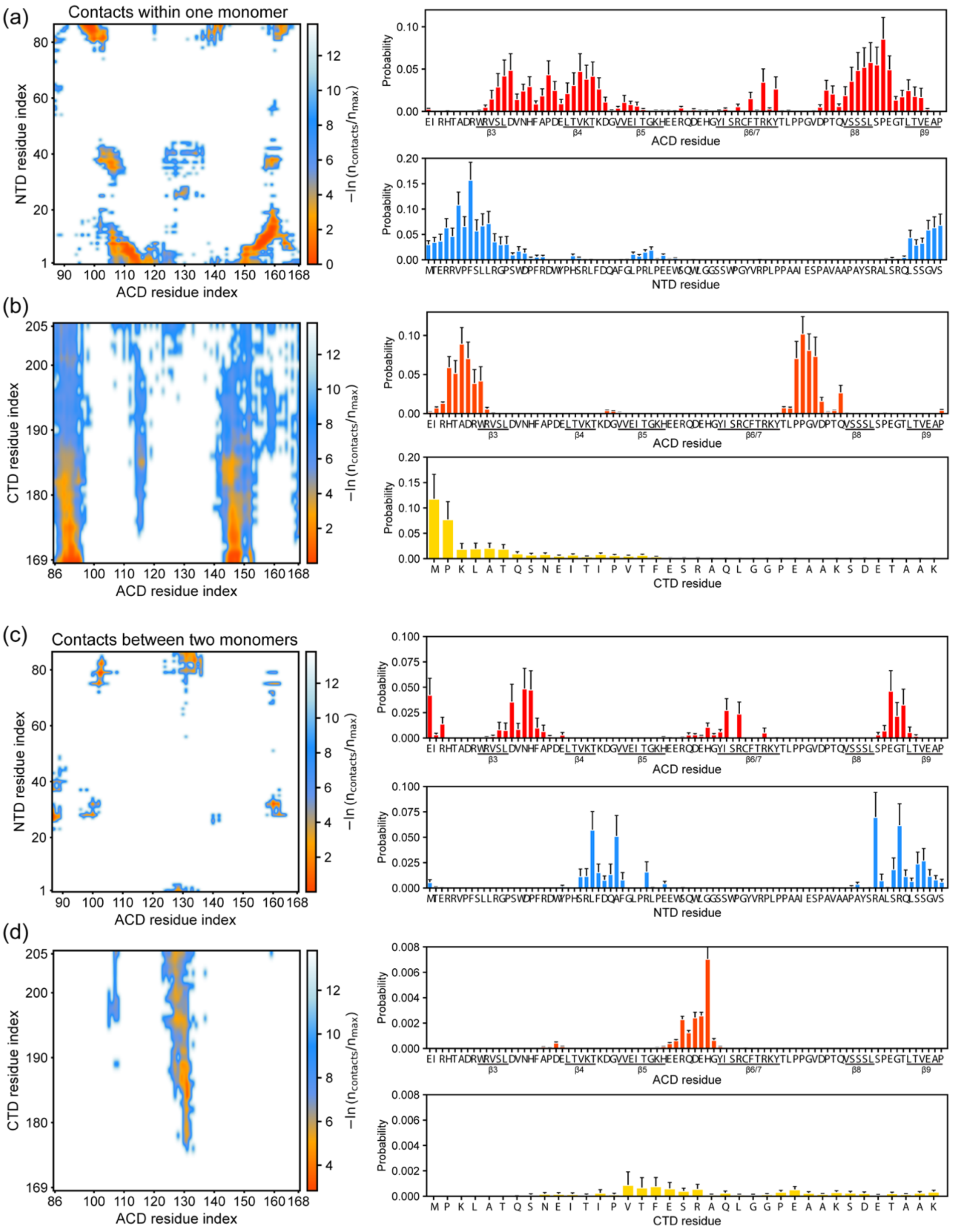


**Figure S15**. Zoomed in contact maps and residue specific contact probabilities for HSPB1 dimers from all-atom molecular dynamics simulations. **(a)** NTD-ACD contacts within each monomeric building block, **(b)** ACD-CTD contacts within each monomeric building block, **(c)** NTD-ACD contacts between monomers, **(b)** ACD-CTD between monomers.

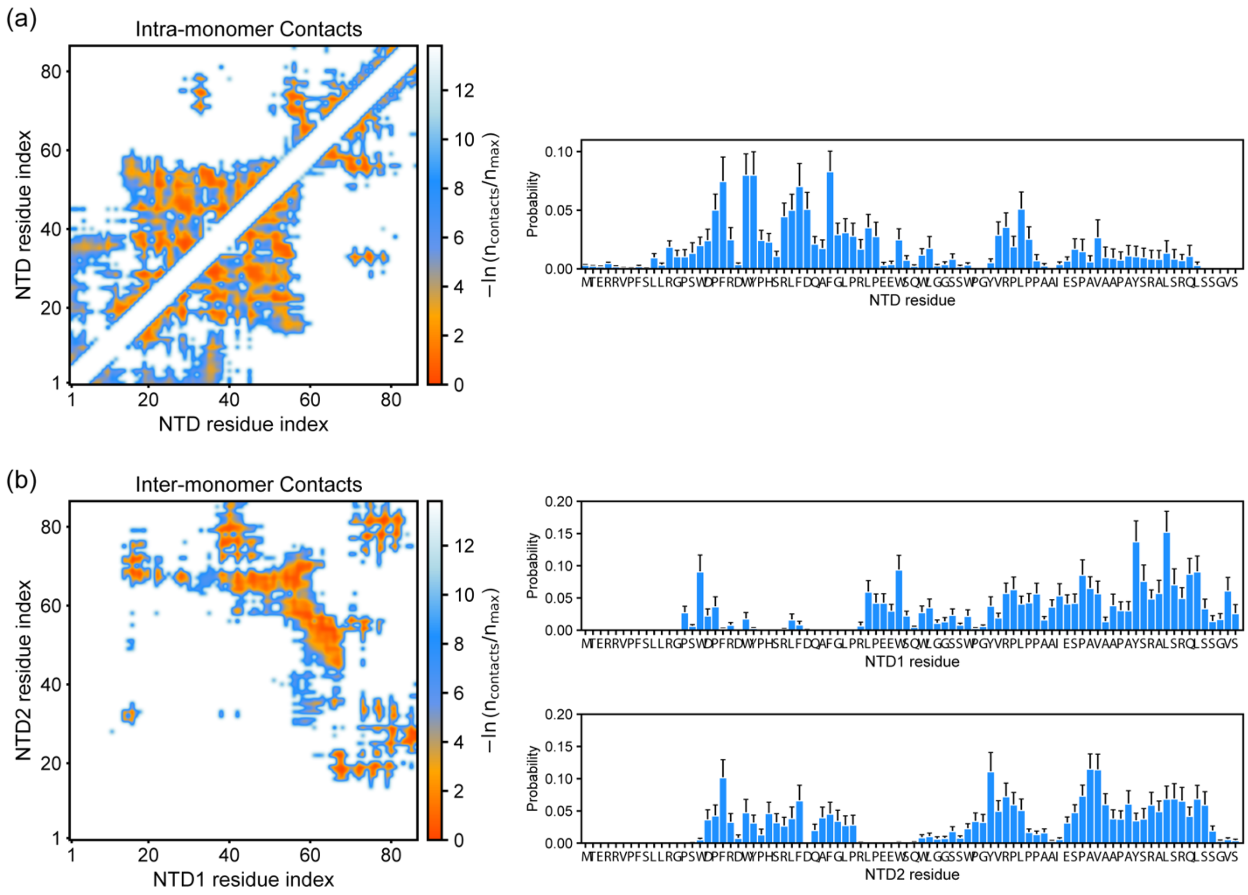


**Figure S16**. Zoomed in contact maps and residue specific contact probabilities for HSPB1 dimers from all-atom molecular dynamics simulations. **(a)** NTD-NTD contacts within each monomeric building block, **(b)** NTD-NTD contacts between monomers.

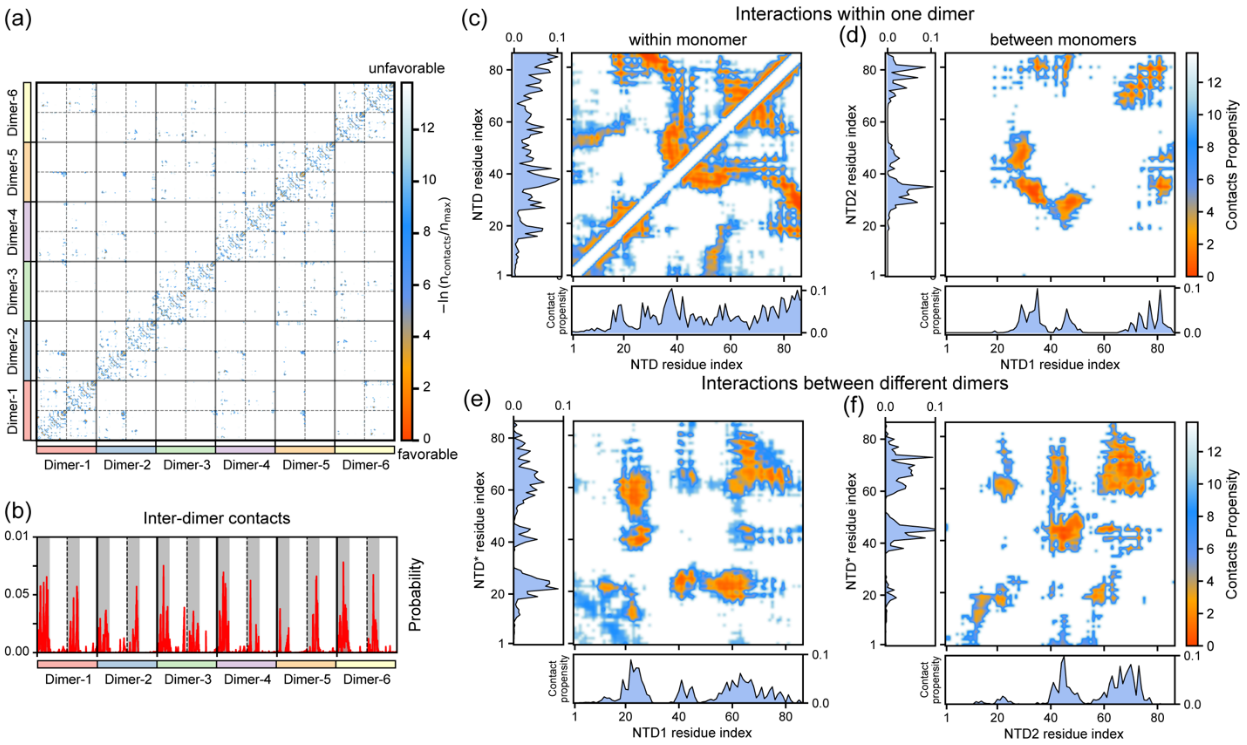


**Figure S17**. Zoomed in contact maps and contact probabilities for HSPB1 dodecamers from all-atom molecular dynamics simulations. **(a)** Intra-dimer (diagonal boxes) and inter-dimer (off-diagonal boxes) contact maps**. (b)** One-dimensional inter-dimer contact map, which was summed over the time-averaged inter-dimer contacts for each residue (off-diagonal blocks). Shaded areas show the NTD regions. Average contact probability within a dimer: **(c)** contacts within a single NTD, and **(d)** contacts between two NTDs in a dimeric building block of the dodecamers. **(e, f)** Average contact probability between the NTD1/NTD2 of a dimer and those (NTD*) of different dimers in the dodecamer.

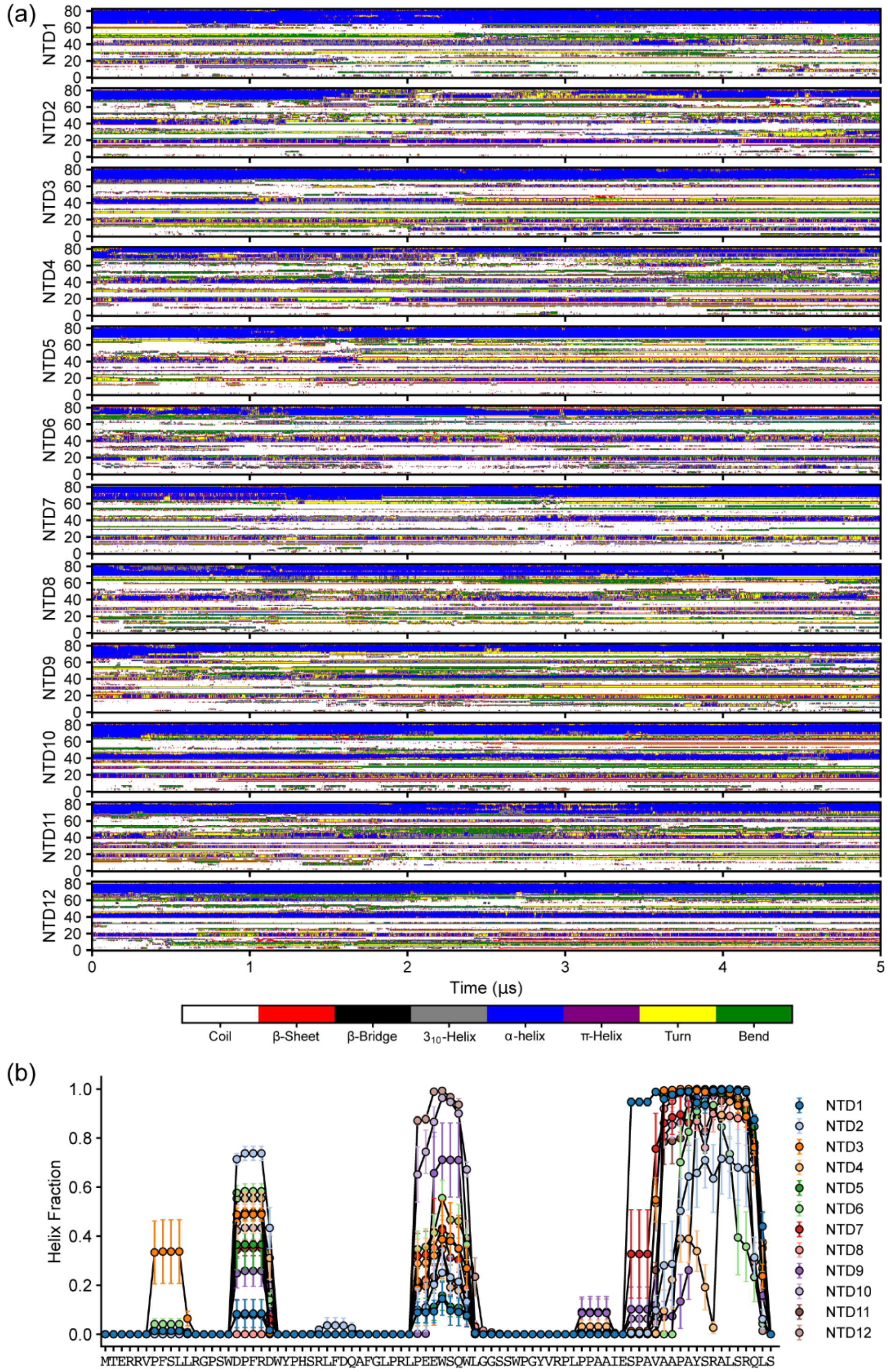


**Figure S18. (a)** Secondary structure evolution as a function of time and **(b)** helix fraction of the NTDs in the HSPB1 dodecamer simulation.

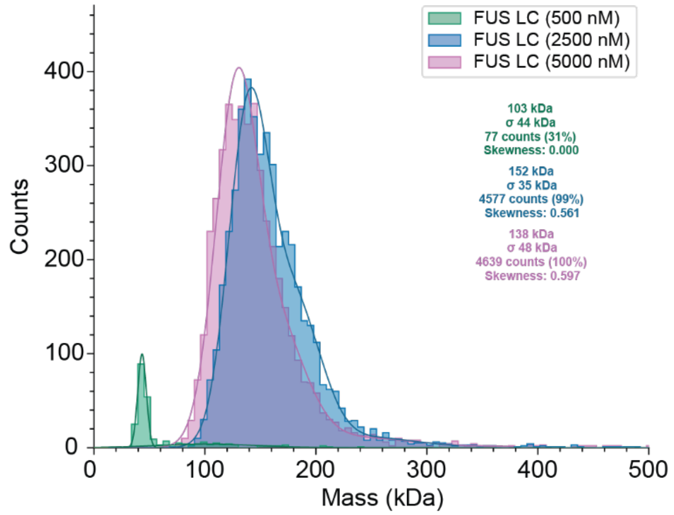


**Figure S19.** Mass photometry gaussian-fit analysis of FUS LC at various protein concentrations. The molecular weight of the FUS LC monomer is 17.225 kDa. Measurements were done in a 150 mM NaCl and 50 mM sodium phosphate (pH=7.2-7.4) buffer.

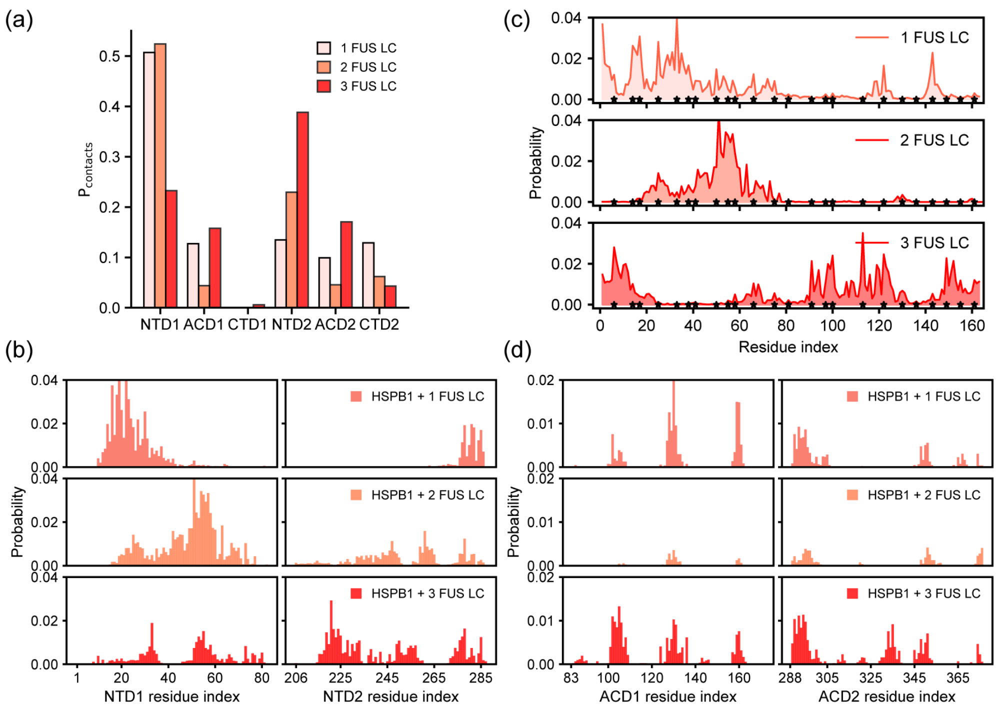


**Figure S20.** One dimensional contact map in the all-atom simulations of an HSPB1 dimer in the presence of one, two, and three FUS LC monomers. **(a)** Summed contact probability for each domain of HSPB1 for interactions with FUS LC. **(b)** Average contact probability per residue between the NTDs and FUS LC. **(c)** Average contact probability per residue of FUS LC for interactions with HSPB1. Tyrosine residues in FUS LC sequence are marked by asterisks (*). **(d)** Average contact probability per residue between ACD1/ACD2 and FUS LC.

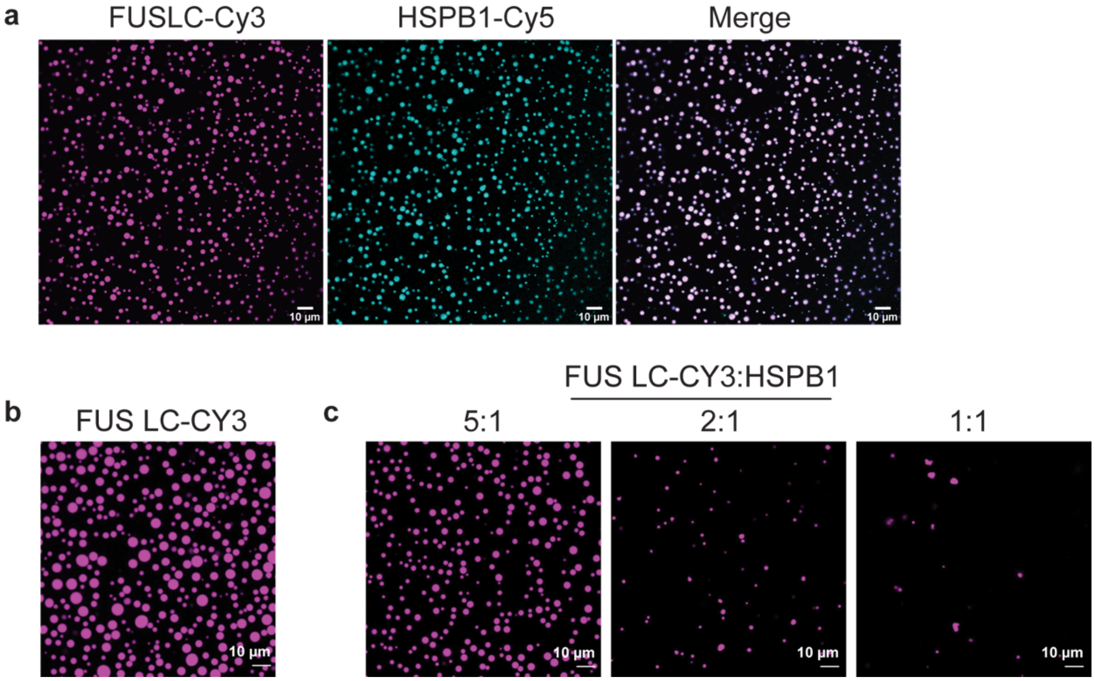


**Figure S21**. **(a)** Fluorescence microscopy images of 5:1 FUS LC (300 µM): HSPB1 (60 µM) condensates to investigate the co-localization of the two proteins. FUS LC was labeled using the Cy3 fluorophore, while HSPB1 was labeled using Cy5. Contrast levels were boosted by using 35% for HSPB1-Cy5 to help with visualization, while for FUS LC 5% labeled protein was sufficient. **(b)** FUS LC condensates (150 µM) without the addition of HSPB1. **(c)** FUS LC condensates (150 µM) in the presence of different ratios of HSPB1. In all cases, FUS LC phase-separation was induced by incubating the protein in 150 mM NaCl and 50 mM sodium phosphate buffer (pH=7.2-7.4) at 4˚C.

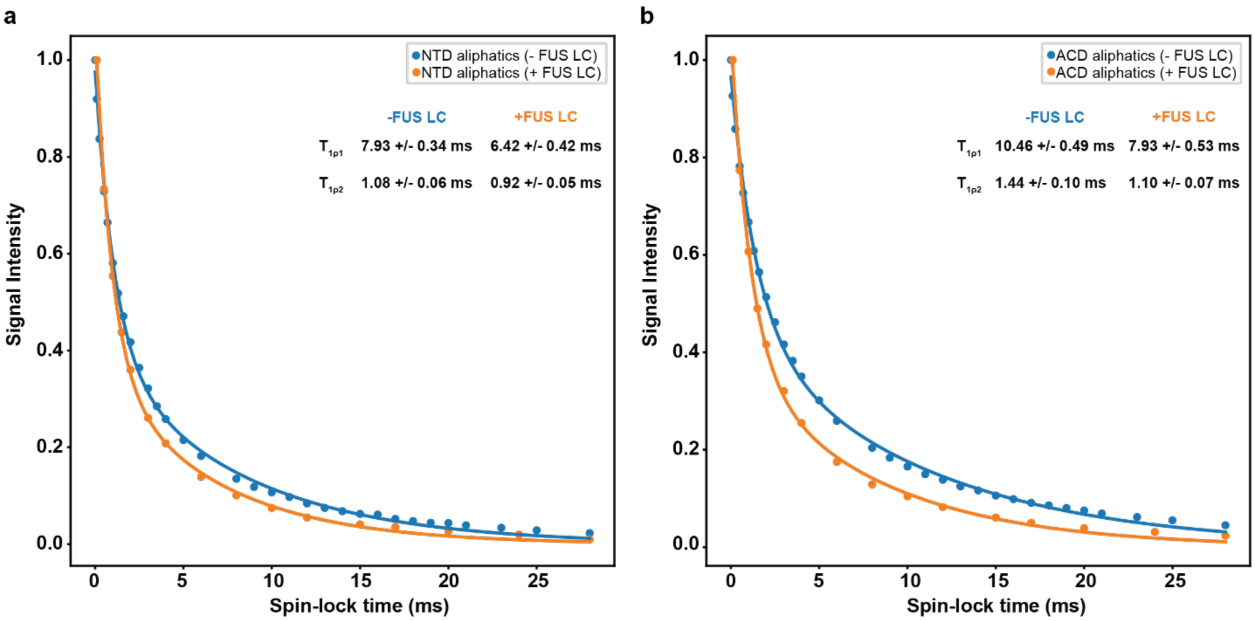


**Figure S22.** Aliphatic $^{13}C$ $T_{1rho}$ measurements of segmentally labeled HSPB1 oligomers on their own and within FUS LC condensates. **(a)** NTD-labeled HSPB1, and **(b)** ACD-CTD labeled HSPB1. A biexponential decay function was used for fitting the experimental data. Data was acquired on a 750 MHz NMR spectrometer with an MAS frequency of 11.111 kHz.

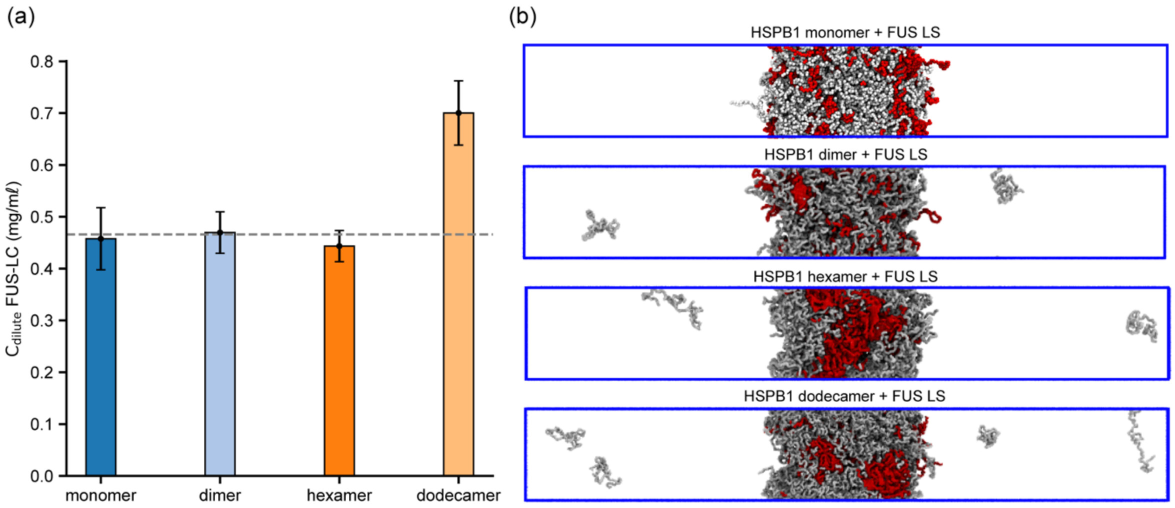


**Figure S23. (a)** Dilute concentrations of FUS LC in coarse-grained co-partitioning simulations with HSPB1 monomers, dimer, hexamers and dodecamers. **(b)** Snapshots of the simulations. FUS LC and HSPB1 are shown in gray and red, respectively. In these simulations, the total concentration was held constant, with the mole fraction of FUS LC to HSPB1 maintained at 2:1.

#### Supporting information movies

**Movie S1**. All-atom (AA) simulation of an HSPB1 monomer.
**Movie S2**. AA simulation of an HSPB1 dimer.
**Movie S3**. Structures of HSPB1 dodecamers predicted without using PDB templates.
**Movie S4**. Structures of HSPB1 dodecamers predicted using PDB templates.
**Movie S5**. AA simulation of an HSPB1 dodecamer.
**Movie S6**. AA simulation of an HSPB1 dimer with 1 FUS LC chain.
**Movie S7**. AA simulation of an HSPB1 dimer with 2 FUS LC chains.
**Movie S8**. AA simulation of an HSPB1 dimer with 3 FUS LC chains.
**Movie S9**. Coarse-grained (CG) coexistence simulations of FUS LC and HSPB1 monomers, dimers, hexamers, and dodecamers.
**Movie S10**. CG coexistence simulations of FUS LC and HSPB1 dimers with various truncations.

#### Supporting information tables

**Table 1.** NMR parameters for routine MAS experiments on fully labeled (FL) HSPB1, and segmentally labeled (SL) NTD and ACD samples. Experiments were performed in a 3.2 mm zirconia rotor (11.111 kHz MAS) on a 750 MHz spectrometer.

| **Sample** | **FL HSPB1** | | **SL HSPB1(NTD)** | | **SL HSPB1(ACD)** | |
|---|---|---|---|---|---|---|
| **Sample Amount** | 30 | mg | 5 | mg | 9 | mg |
| | | | **CP** | | | |
| **Recycle delay** | 5 | s | 5 | s | 5 | s |
| **13C pi/2** | 50 | kHz | 50 | kHz | 50 | kHz |
| **1H pi/2** | 80.6 | kHz | 80.6 | kHz | 80.6 | kHz |
| **CP contact time** | 1000 | μs | 500 | μs | 500 | μs |
| **Decoupling method** | SPINAL64 | | SPINAL64 | | SPINAL64 | |
| **Decoupling power** | 80.6 | kHz | 80.6 | kHz | 80.6 | kHz |
| **Acquisition time** | 9.9 | ms | 10.2 | ms | 10.2 | ms |
| | | | **INEPT** | | | |
| **Recycle delay** | 4 | s | 4 | s | 4 | s |
| **13C pi/2** | 50 | kHz | 50 | kHz | 50 | kHz |
| **1H pi/2** | 80.6 | kHz | 80.6 | kHz | 80.6 | kHz |
| **Refocusing parameter** | 140 | Hz | 140 | Hz | 140 | Hz |
| **Decoupling method** | SPINAL64 | | SPINAL64 | | SPINAL64 | |
| **Decoupling power** | 80.6 | kHz | 80.6 | kHz | 80.6 | kHz |
| **Acquisition time** | 9 | ms | 9 | ms | 9 | ms |
| | | | **DP** | | | |
| **Recycle delay** | 5 | s | 5 | s | 5 | s |
| **13C pi/2** | 50 | kHz | 50 | kHz | 50 | kHz |
| **Decoupling method** | SPINAL64 | | SPINAL64 | | SPINAL64 | |
| **Decoupling power** | 80.6 | W | 80.6 | W | 80.6 | W |
| **Acquisition time** | 10.2 | ms | 10.2 | ms | 10.2 | ms |
| | | | **CP-DARR** | | | |
| **Recycle delay** | 5 | s | 5 | s | 5 | s |
| **13C pi/2** | 50 | kHz | 50 | kHz | 50 | kHz |
| **1H pi/2** | 80.6 | kHz | 80.6 | kHz | 80.6 | kHz |
| **CP contact time** | 400 | μs | 500 | μs | 500 | μs |
| **Decoupling method** | SPINAL64 | | SPINAL64 | | SPINAL64 | |
| **Decoupling power** | 80.6 | W | 80.6 | W | 80.6 | W |
| **DARR mixing time** | 20 | ms | 20 | ms | 20 | ms |
| **1H power during mixing** | 11.11 | kHz | 11.11 | kHz | 11.11 | kHz |

| | | | | | | |
|---|---|---|---|---|---|---|
| **t1 acquisition** | 3.2 | ms | 3.2 | ms | 3.2 | ms |
| **t1 points** | 256 | | 256 | | 256 | |
| | | | **HN-INEPT** | | | |
| **Recycle delay** | 3 | s | | | | |
| **15N pi/2** | 42 | kHz | | | | |
| **1H pi/2** | 79.4 | kHz | | | | |
| **Refocusing paramater** | 92 | Hz | | | | |
| **Decoupling method** | SWf-TPPM | | | | | |
| **Decoupling power** | 79.4 | kHz | | | | |
| **t1 acquisition** | 13 | ms | | | | |
| **t1 points** | 320 | | | | | |
| | | | **HC-INEPT** | | | |
| **Recycle delay** | 5 | s | | | 5 | s |
| **13C pi/2** | 50 | kHz | | | 50 | kHz |
| **1H pi/2** | 83.1 | kHz | | | 80.6 | kHz |
| **Refocusing paramater** | 140 | Hz | | | 140 | Hz |
| **Decoupling method** | SWf-TPPM | | | | SWf-TPPM | |
| **Decoupling power** | 83.1 | W | | | 160 | W |
| **t1 acquisition** | 33.3 | ms | | | 33.3 | ms |
| **t1 points** | 500 | | | | 500 | |
| | | | **CP T1rho** | | | |
| **Number of scans** | | | 1024 | s | 1024 | s |
| **Recycle delay** | | | 2 | s | 2 | s |
| **13C pi/2** | | | 50 | kHz | 50 | kHz |
| **1H pi/2** | | | 80.6 | kHz | 80.6 | kHz |
| **CP contact time** | | | 500 | μs | 500 | μs |
| **Spin lock pulse minimum length** | | | 10 | μs | 10 | μs |
| **Spin lock pulse maximum length** | | | 28000 | μs | 28000 | μs |
| **Spin lock number of points** | | | 32 | | 32 | |
| **Decoupling method** | | | SPINAL64 | | SPINAL64 | |
| **Decoupling power** | | | 80.6 | kHz | 80.6 | kHz |
| **Acquisition time** | | | 9 | ms | 9 | ms |

**Table 2.** NMR parameters for routine MAS experiments on segmentally labeled (SL) NTD and ACD samples in the presence of FUS LC in a phase-separated state. Experiments were performed in a 3.2 mm zirconia rotor (11.111 kHz MAS) on a 750 MHz spectrometer.

| **Sample** | **LLPS SL HSPB1 (NTD)** | | **LLPS SL HSPB1 (ACD)** | |
|---|---|---|---|---|
| **Sample Amount** | 0.5-0.6 | mg | 0.3-0.4 | mg |
| | **CP** | | | |
| **Recycle delay** | 5 | s | 5 | s |
| **13C pi/2** | 50 | kHz | 50 | kHz |
| **1H pi/2** | 80.6 | kHz | 80.6 | kHz |
| **CP contact time** | 500 | μs | 500 | μs |
| **Decoupling method** | SPINAL64 | | SPINAL64 | |
| **Decoupling power** | 80.6 | kHz | 80.6 | kHz |
| **Acquisition time** | 10.2 | ms | 10.2 | ms |
| | **INEPT** | | | |
| **Recycle delay** | 4 | s | 4 | s |
| **13C pi/2** | 50 | kHz | 50 | kHz |
| **1H pi/2** | 80.6 | kHz | 80.6 | kHz |
| **Refocusing parameter** | 140 | Hz | 140 | Hz |
| **Decoupling method** | SPINAL64 | | SPINAL64 | |
| **Decoupling power** | 80.6 | kHz | 80.6 | kHz |
| **Acquisition time** | 9 | ms | 9 | ms |
| | **DP** | | | |
| **Recycle delay** | 5 | s | 5 | s |
| **13C pi/2** | 50 | kHz | 50 | kHz |
| **Decoupling method** | SPINAL64 | | SPINAL64 | |
| **Decoupling power** | 80.6 | W | 80.6 | W |
| **Acquisition time** | 10.2 | ms | 10.2 | ms |
| | **CP T1rho** | | | |
| **Number of scans** | 10240 | s | 10240 | s |
| **Recycle delay** | 2 | s | 2 | s |
| **13C pi/2** | 50 | kHz | 50 | kHz |
| **1H pi/2** | 80.6 | kHz | 80.6 | kHz |
| **CP contact time** | 500 | μs | 500 | μs |
| **Spin lock pulse minimum length** | 100 | μs | 100 | μs |
| **Spin lock pulse maximum length** | 28000 | μs | 28000 | μs |
| **Spin lock number of points** | 16 | | 16 | |
| **Decoupling method** | SPINAL64 | | SPINAL64 | |
| **Decoupling power** | 80.6 | kHz | 80.6 | kHz |
| **Acquisition time** | 9 | ms | 9 | ms |

**Table 3.** NMR parameters for fast $^{1}H$-based MAS experiments. Experiments were performed in a 0.4 mm zirconia rotor (160 kHz MAS) on an 800 MHz NMR spectrometer, packed with approximately 25 µg of HSPB1(NTD-[$^{13}C$,$^{15}N$-ACD-CTD]).

| Experiment | CP-hNH | | | CP-hCH | |
|---|---|---|---|---|---|
| **Number of scans** | 1280 | | **Number of scans** | 112 | |
| **Recycle delay** | 1.8 | s | **Recycle delay** | 1.5 | s |
| **1H pi/2** | 233.6 | kHz | **1H pi/2** | 233.6 | kHz |
| **15N pi/2** | 39.06 | kHz | **13C pi/2** | 73.5 | kHz |
| **CP contact time (HtoN)** | 1000 | µs | **CP contact time (HtoC)** | 800 | µs |
| **CP contact time (NtoH)** | 800 | µs | **CP contact time (CtoH)** | 400 | µs |
| **Decoupling method** | waltz16 | | **Decoupling method** | waltz16 | |
| **Decoupling power (1H)** | 0.0254 | W | **Decoupling power (1H)** | 0.217 | W |
| **Decoupling power (15N)** | 0.167 | W | **Decoupling power (13C)** | 0.185 | W |
| **t1 acquisition** | 3.2 | ms | **t1 acquisition** | 10.8 | ms |
| **t1 points** | 256 | | **t1 points** | 800 | |
| **t2 acquisition** | 9 | ms | **t2 acquisition** | 20 | ms |
| **t2 points** | 1024 | | **t2 points** | 952 | |
